# PCNA-Polκ-Polδ/USP18 axes stabilize replication fork and restart to reduce cisplatin cytotoxicity

**DOI:** 10.64898/2026.06.08.730955

**Authors:** Ipsita Subhadarsini, Jugal Kishor Sahu, Shweta Thakur, Rupesh Dash, Narottam Acharya

**Affiliations:** Laboratory of Genomic Instability and Diseases, Department of Infectious Disease Biology, Institute of Life Sciences, Bhubaneswar-751023, India; Regional Center of Biotechnology, Faridabad, India

**Keywords:** Translesion DNA synthesis, chemoresistance, DNA Replication, PCNA, γH2AX, Apoptosis, Homologous Recombination, checkpoints, ubiquitin, proteasome, USP18

## Abstract

Cisplatin and its analogues are valuable anti-cancer drugs that target the genome, block DNA replication, and induce apoptosis. As a counteractive response, cancer cells activate several mechanisms to maintain uninterrupted DNA replication, and these are yet to be fully elucidated. This study using head and neck squamous cell carcinoma (HNSCC) demonstrated the involvement of DNA polymerase Kappa (Polκ), a translesion DNA synthesis (TLS) polymerase that primarily functions as a mismatch extender, in cisplatin resistance. Interestingly, the catalytic activity of Polκ plays a minimal role in adduct bypass; rather, tripartite interactions involving it rewire and stabilize the stalled replication fork. While the Polκ-PCNA-Polδ axis facilitates efficient proliferation of cisplatin-resistant cells, the Polκ-PCNA-USP18 axis stabilizes critical proteins of the ATM-ATR, and HR and NHEJ pathways to protect the replication fork, repair damage, and restart DNA synthesis under cisplatin-induced stress. In resistant cells, the efficiency of ubiquitin-mediated proteasomal degradation is low and is further diminished by PCNA-bound Polκ-recruited USP18 deubiquitinase, thereby maintaining cellular homeostasis. In conclusion, for the first time, we uncovered two critical Polκ axes crucial for regulating cisplatin toxicity in cells and provided the foundation for future drug discovery against advanced HNSCC by targeting this non-essential DNA polymerase.

## Introduction

Cisplatin (cis-diamminedichloroplatinum (II)) is one of the widely used metal-based chemotherapeutic drugs to treat cancers like testicular, ovarian, bladder, lung, cervical, head and neck, gastric cancers, etc. (Dasari and Tchounwou 2014; Ghosh 2019). Cisplatin exerts its anticancer activity by generating Pt-DNA adducts by cross-linking the N7 atoms of adjacent purines preferably guanines on the same or different DNA strands, at multiple sites across the nuclear and mitochondrial genomes simultaneously. Thus, the generated lesions block both DNA replication and transcription and activate several transduction pathways, ultimately leading to apoptosis in rapidly proliferating tumor cells. The clinical success of cisplatin is compromised by the development of intrinsic and acquired chemoresistance, as well as dose-limiting toxicity. Reduced efficacy of cisplatin depends on multiple factors, such as reduced drug accumulation by efficient efflux pumps and sequestration by certain proteins, inactivation of the drug by binding with different proteins, success in inducing DNA damage, increased cellular DNA repair ability, activation of DNA damage tolerance pathways, and alteration of different proteins that signal to apoptosis (Maji et al. 2018).

Repair and bypass of cisplatin DNA adducts by nucleotide excision repair (NER) and by specialized DNA polymerases during translesion DNA synthesis (TLS), respectively, are key to cisplatin resistance (Duan et al. 2020). Studies have reported that low intrinsic expression of NER proteins and low NER activity are the main reasons for the impressive clinical success of cisplatin against metastatic testicular germ cell tumors (Masters and Koberle 2003; Welsh et al. 2004). Recent CRISPR-Cas9 based screening analysis suggested a role of transcription-coupled NER (TC-NER) in cisplatin resistance (Slyskova et al. 2018). Interstrand cisplatin cross-links are easily recognized and repaired by NER, whereas less efficient recognition of intrastrand cross-links leads to their higher abundance in cells and, therefore, they are highly cytotoxic (Sarkar et al. 2006). TLS is an alternative DNA replication process that enables bypass of DNA lesions encountered during replication, which are usually not handled by high fidelity replicative DNA polymerases Polδ and Polε, and is emerging as a primary chemotherapeutic target (Acharya et al. 2020a). Not only TLS polymerases protect normal cells from cisplatin-induced genome instability during chemotherapy, but they also reduce cisplatin toxicity in tumor cells by preventing replication fork collapse and cell death, and thereby contributing to cisplatin resistance. Human cells harbour Y-family DNA polymerases such as REV1, Pol17, Polτ, and Polκ that bypass specific DNA lesions (Acharya et al. 2020a). Among these, the role of Polη in bypassing Pt-GG lesions has been extensively studied both in yeast and humans (Ummat et al. 2012; Zhao et al. 2012; Manohar et al. 2018). An increased expression of Polη has impacted cisplatin therapy and survival time of patients with non–small-cell lung cancer, ovarian cancer, and metastatic gastric adenocarcinoma (Chin et al. 2008; Srivastava et al. 2015).

Co-crystal structural analyses suggested that the catalytic pocket of Pol17 accommodates the Pt-GG lesion without steric hindrance and enables successful error-free bypass (Ummat et al. 2012; Zhao et al. 2012). Conversely, an alternate Polι/Polθ-dependent error-prone TLS pathway can replicate through cisplatin-induced intrastrand cross-links. For both pathways, Rev1 is an indispensable scaffolding component of Pol17 and Polτ (Lin et al. 2006; Yoon et al. 2015; Yoon et al. 2021). Although Polσ, a complex of Rev3 and Rev7, belongs to the B-family, it can participate in TLS (Baynton et al. 1998). While Polσ is not required for TLS opposite cisplatin-induced intrastrand cross-links in normal cells, studies suggest that cancer cells require Rev1-Polζ, and depletion of Rev1 or Rev3 sensitizes cancer cells to cisplatin, indicating a role for Rev1-Polζ in chemoresistance (Doles et al. 2010; Zhu et al. 2016). Another study revealed that Pol17 contributes minimally in promoting TLS through cisplatin analogues like oxaliplatin, satraplatin, and picoplatin, whereas Rev1 and Polσ play a critical role in bypassing all four platinum analogs (Sharma et al. 2012). In contrast to other TLS Pols, the role of Polκ in lesion bypass remains poorly defined. Biochemical assays suggested that Polκ is largely inefficient at inserting nucleotides opposite cisplatin-induced major groove DNA lesions; however, it can act as a mismatch extender to facilitate complete bypass (Haracska et al. 2002; Wolfle et al. 2003; Vasquez-Del Carpio et al. 2011; Jha and Ling 2018). Polκ can bypass minor groove DNA damage induced by benzo[a]pyrene and acylfuvenes (Ohashi et al. 2000; Ogi et al. 2002). A recent study reported that while bypassing minor groove DNA lesions, Polκ activity is dependent on PCNA ubiquitination but independent of Rev1, whereas in TLS through major groove adducts and abasic sites, Polκ mostly plays a structural role and stabilizes the Rev1–Polζ extension complex on DNA, which is also regulated by ubiquitinated PCNA (Selles-Baiget et al. 2025).

Uninterrupted DNA replication in cancer cells is a major driver of chemoresistance, as tumor cells often develop mechanisms to maintain replication fork progression despite chemotherapy-induced damage, leading to treatment failure. Since we observed overexpression of Polκ in specific tumors upon cisplatin exposure, and among the TLS pols, its role in chemoresistance is the least reported, this study established and explored underlying TLS and non-TLS mechanisms by which Polκ maintains DNA replication continuity to induce cisplatin chemoresistance. We uncovered two axes through which Polκ facilitates uninterrupted DNA replication in cancer cells, thereby nullifying cisplatin toxicity. We found that Polκ rewires replication machinery specifically in cisplatin-resistant cancer cells by establishing a tripartite interaction with PCNA-Polδ to enhance cellular proliferation. PCNA-bound Polκ maintains cellular homeostasis of several checkpoint and DDR proteins by preventing their ubiquitin-mediated proteasomal degradation by recruiting USP18 deubiquitinase to its ubiquitin-binding Zn finger domain (UBZ). Finally, we propose that targeting Polκ and USP18 can enhance the efficacy of cisplatin, a first-line anticancer drug.

## Results

### Polκ is markedly upregulated in specific cancer cells upon cisplatin treatment and in cisplatin-resistant oral squamous cell carcinoma (OSCC)

The expression of DNA polymerase involved in TLS at a specific lesion is usually upregulated when cells are exposed to genotoxic agents that generate such lesions in the genome. To determine a possible role of Polκ in cisplatin resistance, a panel of tumour specific cell lines from breast (MCF7 and MDA-MB-231), brain (SH-SY5Y), liver (Huh7), head and neck (H357 and SCC9), pancreatic (MIAPaCa-2), prostate (DU145) and lungs (A549) cancers along with a transformed but non-tumorigenic cell line HEK293 (Human embryonic kidney) were treated with 10 μM of cisplatin for 24 hrs and the expression of Polκ was determined by Western analysis (**Figure 1 A and B**). Since the role of Pol17 in cisplatin chemoresistance has already been reported, we also checked its expression in these cells. We observed a tumour-specific induced expression of Pol17 and Polκ upon cisplatin exposure. In lung cancer and noncancerous HEK293 cells, Pol17 expression increased 2-3-fold upon cisplatin treatment compared with untreated controls; however, no significant change in its expression was observed in other cell types. Polκ showed induced expression in HNSCC cell lines treated with cisplatin, but not in other tumour cell lines. Interestingly, the pancreatic cancer cells showed more expression of both Pol17 and Polκ upon cisplatin exposure. This result suggested that cisplatin resistance in a particular tumour could be Pol17 or Polκ DNA polymerase-specific.

**Figure 1:**
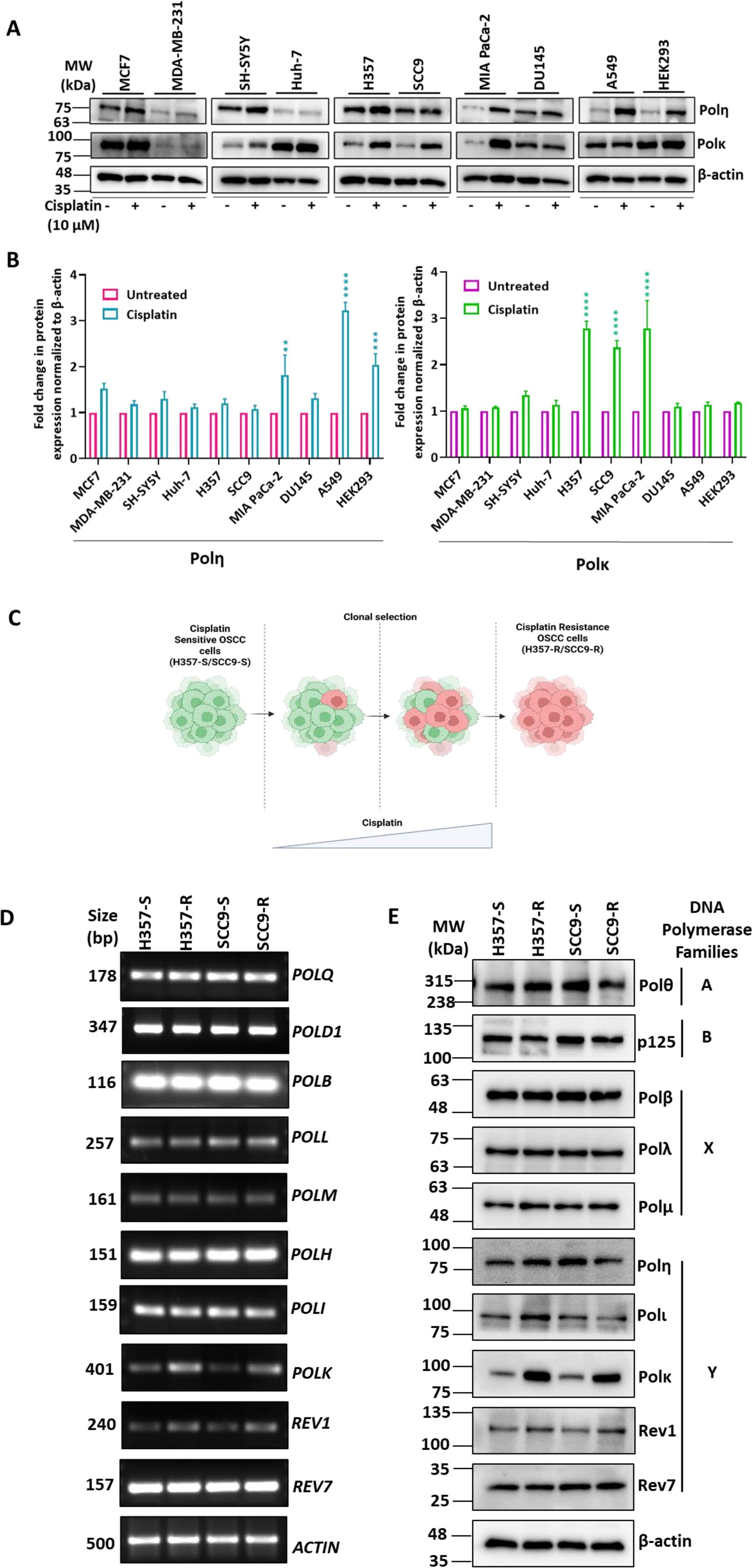
Dysregulation of Polκ in cisplatin treated and resistant cancer cells. **(A)** Western blot analyses of various cell-free lysates from cell lines, untreated and treated with 10 μM cisplatin, were performed using Polη and Polκ antibodies. β-actin was used as a loading control. **(B)** Protein band intensities were estimated, and fold change after β-actin normalization was shown in a bar graph as mean ± SEM (n=2). Statistical significance determined by two-way ANOVA, Šídák’s multiple comparisons test (**p < 0.01, ***p < 0.001, ****p < 0.0001). **(C)** Schematic representation of generation of cisplatin resistant OSCC cell lines (H357-R and SCC9-R) from their respective cisplatin sensitive H357-S and SCC9-S cells. **(D)** RT-PCR analysis of various DNA polymerases in cisplatin resistant and sensitive OSCC cells. ACTIN was taken as a loading control. **(E)** Western blot analyses of lysates of OSCC cells using specific antibodies against various DNA polymerases or their subunits as mentioned. β-actin was detected as a loading control.

To further strengthen this, we assessed and compared the expression of various DNA polymerases in cisplatin-resistant H357 (H357-R) and SCC9 (SCC9-R) cells with that in their sensitive parental isogenic counterparts (H357-S and SCC9-S). H357-R and SCC9-R cells have been used in several studies and can tolerate up to 40 µM cisplatin, which is 3-4-fold higher than their IC50 values (10-15 µM) (Mohapatra et al. 2021; Shriwas et al. 2021). Briefly, these resistant cells were generated by chronic exposure to a gradually increasing dose of 10 µM cisplatin over 8 months, and resistant cells were selected (**Figure 1C**). Human cells possess 15 DNA polymerases that belong to A, B, X, and Y families (Acharya et al. 2020a). The expression in terms of transcripts and protein levels of most of these Pols were checked by both reverse transcription PCR (RT-PCR) and Western analyses (**Figure 1 D and E, Supp. Figure 1A i and ii**). p125 and Rev7 were checked as a subunit of Polδ (p125-p50-p68-p12) and Polσ (Rev3-Rev7) holoenzyme complexes, respectively. In these cisplatin resistant cells, the transcript and protein levels of most of these polymerases did not alter significantly except Rev1 and Polκ. To our surprise, notable polymerases such as Pol17 and Polσ, which have been reported to contribute to cisplatin resistance across various cancer types, remained unaltered in resistant OSCC cells. Since Polκ was more highly dysregulated in resistant cells than Rev1, its role in chemoresistance was further explored in greater detail.

### Polκ depletion re-sensitizes resistant OSCC cells to cisplatin

To investigate how *POLK* deficiency would affect cisplatin sensitivity in resistant cancer cells, we decided to delete the gene using the CRISPR-Cas9 technique in H357-R and SCC9-R cells. While we were able to successfully generate and select *POLK*^-/-^ clones of H357-R (H357-R *POLK* KO), as confirmed by Western blotting, despite our several attempts, we could not generate a similar clone in SCC9 **(Supp. Figure 1B and Figure 2A lanes 5 and 6)**. Therefore, for subsequent analyses, H357-R *POLK* KO cells were considered. When the H357-S cells were subjected to cisplatin treatment, a 2 to 3 fold increase in Polκ expression was observed in comparison to untreated cells; however, as shown in Fig. 1 D, the Polκ level was higher in H357-R than in H357-S irrespective of cisplatin presence or absence (**Fig. 2A, compare lane 1 with 2-4, Supp. Figure 1C**). This result was further confirmed by immunofluorescence analysis, where a higher amount of immune-stained Polκ was observed in resistant than sensitive cells **(Supp. Figure 1D i and ii)**. MTT-based and clonogenic cell survival assays revealed that while H357-R cells were resistant to cisplatin, both H357-S and H357-R *POLK* KO cells were hypersensitive, *albeit* more so in the Polκ-deficient cells (**Fig. 2 B and C**). IC50 values of H357-S and H357-R *POLK* KO cells were nearly 6-10µM, whereas for H357-R, it was estimated to be >32µM, which is consistent with the earlier published report. Cell death due to cisplatin treatment was assessed by Annexin-V and PI staining and FACS analyses, which further confirmed that the depletion of Polκ re-sensitizes OSCC resistant cells to cisplatin, as a higher percentage of apoptotic cells was observed in both H357-S and H357-R *POLK* KO cells upon cisplatin treatment (**Fig. 2D i and ii**). As expected, a higher percentage of the resistant H357-R cells still survived despite cisplatin treatment. Altogether, our results confirmed that overexpressed Polκ induces cisplatin chemoresistance in H357 OSCC cells. To further demonstrate the role of *POLK* in cisplatin resistance, *POLK* gene expression was depleted using specific siRNA in both H357-S and SCC9-S cells, and their sensitivity to cisplatin was assessed (**Supp. Figure 1E and F**). Interestingly, despite the presence of other TLS DNA polymerases, depletion of Polκ alone sensitized these OSCC cells to cisplatin, suggesting its critical role in chemoresistance.

**Figure 2:**
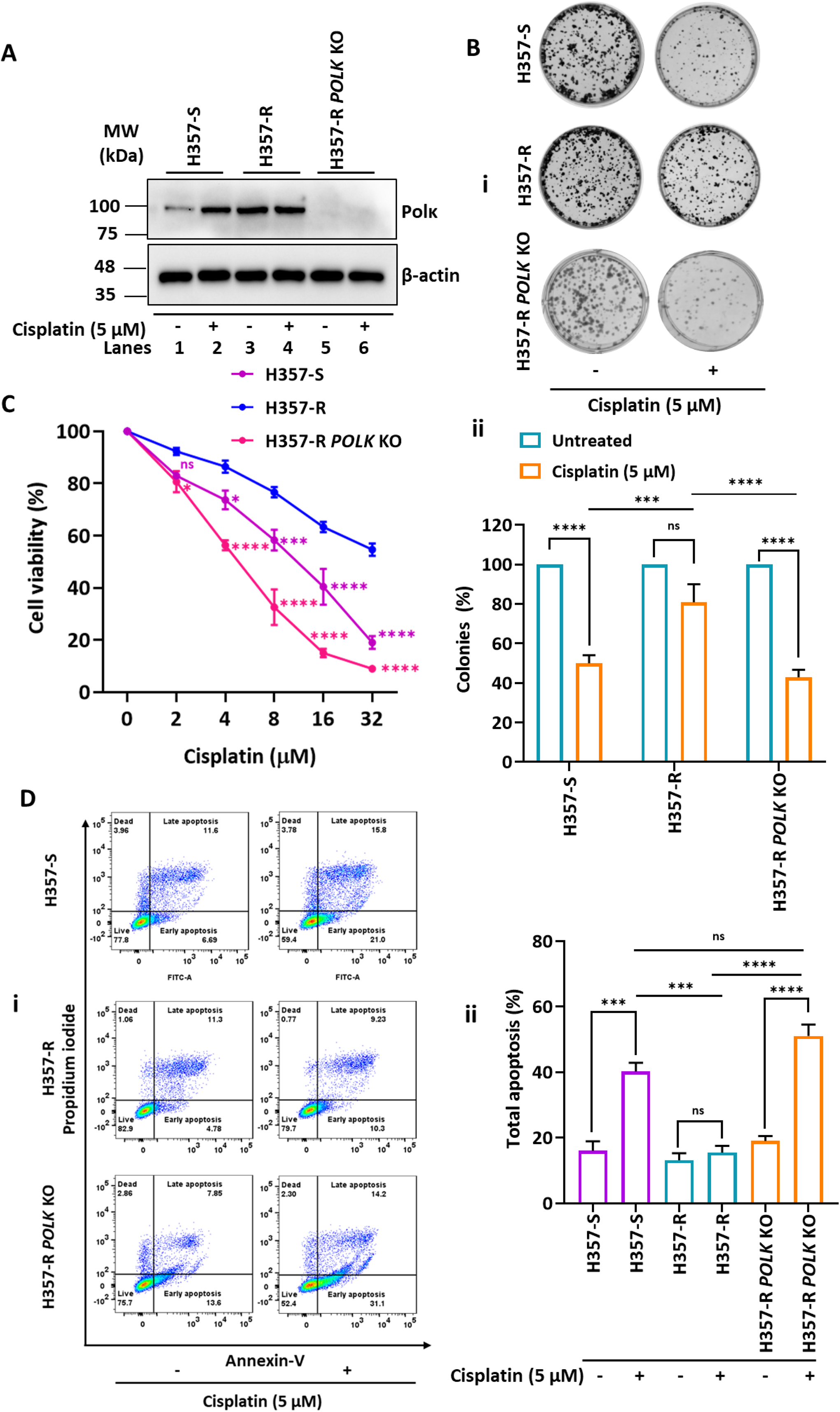
Absence of Polκ re-sensitizes OSCC resistant cells to cisplatin. **(A)** H357-S, H357-R and H357-R *POLK* KO cells were treated with 5 μM cisplatin and analyzed for Polκ expression by Western blot analysis. β-actin was used as a loading control. **(B)** A colony-forming assay was performed using H357-S, H357-R and *POLK* KO cells without and with 5 μM cisplatin (i). Bar graph represents colony percentage as mean ± SEM (n=3) as determined by two-way ANOVA with Tukey’s multiple comparisons test (***p < 0.001, ****p < 0.0001, and ns-nonsignificant) (ii). (**C)** Cells were treated with cisplatin for 48 hours, and then MTT assays were performed. Graph indicates the mean ± SEM (n=3), two-way ANOVA with Tukey’s multiple comparisons test (*p < 0.05, ***p < 0.001, ****p < 0.0001, and ns-nonsignificant). Purple asterisks indicate comparison between H357-R and H357-S; pink asterisks indicate comparison between H357-R and H357-R *POLK* KO. **(D)** After treatment with 5 µM cisplatin for 48 hours, cells were stained with Annexin V/PI and analyzed by flow cytometry. The percentage of cells in each quadrant is as indicated (i). The bar graph represents the percentage of total apoptosis (early and late apoptosis). Data were represented as mean ± SEM (n=3). Statistical significance determined by two-way ANOVA, Tukey’s multiple comparisons test, (***p < 0.001, ****p < 0.0001, and ns-nonsignificant) (ii).

### Polκ depletion enhances genomic instability and delays cell cycle progression in resistant OSCC cells

Since Polκ is a TLS polymerase, its absence can lead to accumulation of damaged DNA from cisplatin adducts associated with replication fork collapse. Therefore, to correlate cellular sensitivity with DNA damage accumulation in Polκ depleted cells, the DNA damage status of these cells following cisplatin treatment was determined. The comet assay confirmed increased DNA tailing of genomic DNA in Polκ-depleted cells following cisplatin treatment, indicating a higher percentage of DNA breaks. While minimal tailing was observed in H357-R cells, H357-S cells also showed DNA tailing when exposed to sub-optimal concentrations of cisplatin (**Figure 3A i and ii**). To further confirm the re-sensitization of H357-R cells to cisplatin due to the lack of Polκ, other DNA damage markers like Gamma-H2A.X (ψH2A.X) and cleaved PARP1 were checked. γH2A.X is a biomarker for the presence of DNA double-strand breaks (DSBs). After a DSB occurs, ATM and DNA-PK phosphorylate the serine-139 residue of histone variant H2A.X, forming γH2A.X foci that act as “anchors” to recruit DNA repair proteins to the site of damage (Mah et al. 2010). Both immunofluorescence and Western analyses suggested that there was a higher percentage of ψH2A.X foci as a cellular response to DNA damage in H357-S and H357-R *POLK* KO than in H357-R cells upon cisplatin treatment (**Figure 3B i and ii, Figure 3C, and Supp. Figure 2A**). The phosphorylation of H2A.X to γH2AX creates binding sites that enhance PARP1’s affinity and its catalytic efficiency, thereby stabilizing PARP1 at the damage site and facilitating DNA repair. However, under conditions of excessive DNA damage, the cell may initiate apoptotic pathways, leading to PARP1 cleavage by activated caspases, producing 89-kDa and 55-kDa fragments (Kumari et al. 2025). We observed more accumulation of cleaved PARP1 when H357-S and H357-R *POLK* KO cells were treated with cisplatin (**Figure 3D, Supp. Figure 2B**). Despite cisplatin treatment, H357-R cells did not show any increase in ψH2A.X foci and PARP1 cleavage, suggesting efficient repair of DNA breaks. Notably, cleaved PARP1 product was observed even in untreated H357-R *POLK* KO cells. These results indicated that, in the absence of *POLK*, cells can accumulate DSBs in the presence of cisplatin. In that context, Polκ depletion may affect cell cycle progression. Therefore, a synchronized population of various cells post a double-thymidine block was allowed to proliferate without and with 5 µM cisplatin, and cells were harvested and analyzed by FACS (**Figure 3E i and ii, Supp. Figure 2C i and ii**). Without cisplatin, both H357-S and H357-R cells progressed through a complete cell cycle within 12 hrs; however, we observed a significant delay in the progression of Polκ-depleted cells, and the majority of the cells were stuck in the G2/M phase (9-12 hours). This observation is corroborated by more ψH2A.X foci and cleaved PARP1 accumulation in H357-R *POLK* KO cells, which were further heightened upon cisplatin treatment. Chronic exposure to cisplatin significantly delayed cell-cycle progression in H357-S and H357-R *POLK* KO cells compared with H357-R cells. At 6 hrs, most of the cells of H357-S and H357-R *POLK* KO were still in G1/S phase, while H357-R cells were accumulated in G2/M phase (**Supp. Figure 2C**). Since cell cycle progression was affected by *POLK* depletion even in the absence of genotoxic stress, we assessed cellular proliferation by measuring Ki-67 expression. A higher percentage of Ki-67-positive cells usually indicates increased cellular proliferation, and the immunofluorescence assay suggested a higher percentage of Ki-67-positive resistant OSCC cells (85-95%), which decreased significantly upon Polκ depletion (**Supp. Figure 2D i and ii)**. This suggests that Polκ contributes to the proliferative efficacy of H357-R cells. Altogether, these data suggest that Polκ’s function is associated with multiple phenotypes in resistant OSCC cells, as its absence accumulates DNA breaks, induces apoptosis, delays cell cycle progression, and reduces cellular proliferation, both in the absence and presence of cisplatin. Thus, Polκ plays a crucial role in stabilizing and recovering replication forks after cisplatin-induced DNA damage and probably also in replicating undamaged DNA in H357-R cells.

**Figure 3:**
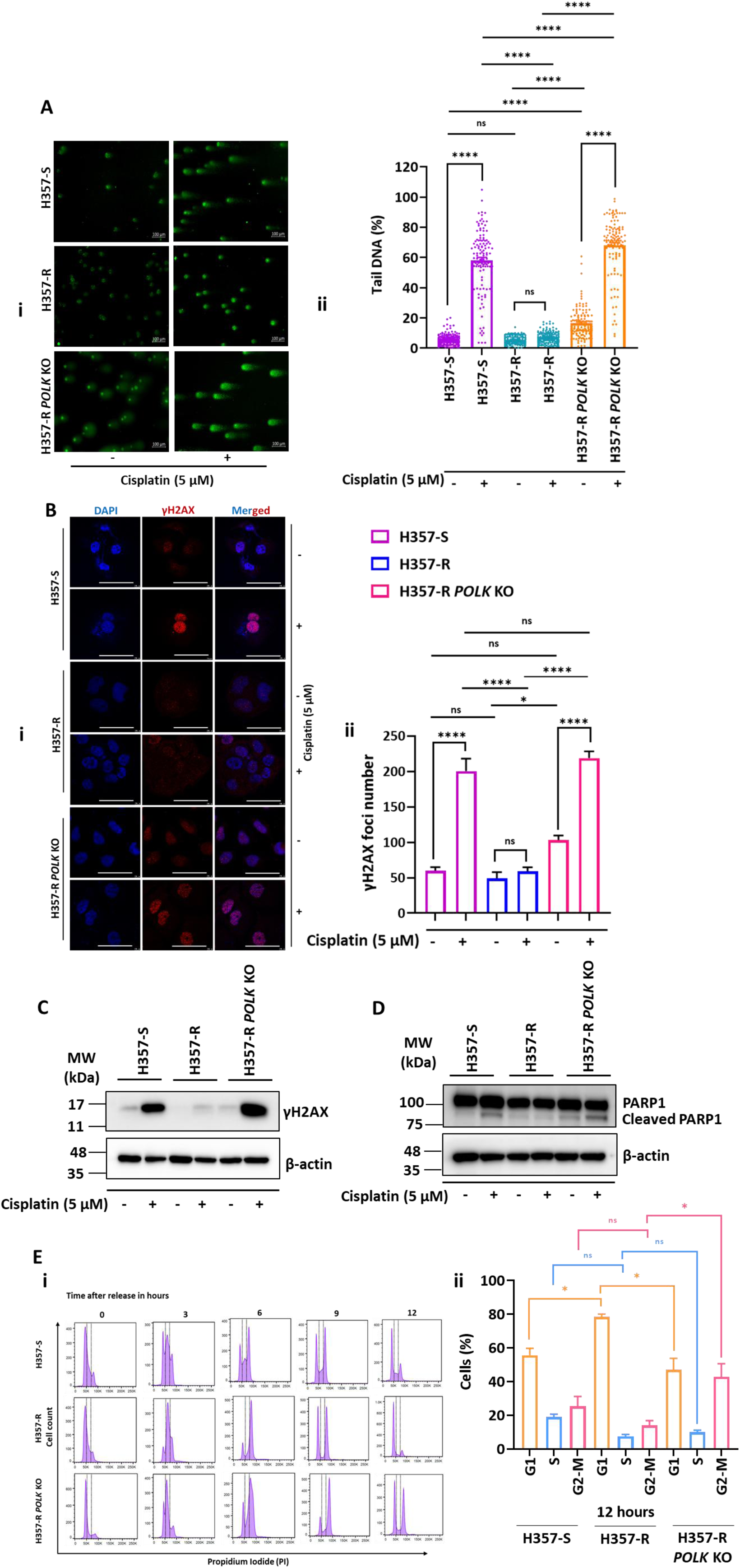
Absence of Polκ enhances genomic instability and delays cell-cycle progression in OSCC-resistant cells. **(A)** Representative alkaline comet images showing DNA tailing in cells following treatment with 5 μM cisplatin for 48 hours (i). The bar graph represents the percentage of tail DNA. Data are represented as (mean ± SEM; n=3). Statistical significance determined by two-way ANOVA with Tukey’s multiple comparisons test (****p < 0.0001, and ns-nonsignificant) (ii). **(B)** Representative immunofluorescence images of H357-S, H357-R and H357-R *POLK* KO cells showing γH2AX (red) puncta and DAPI (blue) and corresponding merged images (i). Quantification of γH2AX foci numbers is indicated in a bar diagram as mean ± SEM (n=3) (ii). Statistical significance determined by Ordinary one-way ANOVA, Tukey’s multiple comparisons test, (*p < 0.05, ****p < 0.0001, and ns-nonsignificant). Scale bars indicate 7.5 μm. **(C)** Western blot analyses of H357-S, H357-R and H357-R *POLK* KO lysate using γH2AX antibody was carried out. β-actin was used as a loading control. **(D)** Western blot analyses of H357-S, H357-R and H357-R *POLK* KO lysates probed with PARP1 antibody were carried out. β-actin was used as a loading control. **(E)** A representative cell cycle profile of H357-S, H357-R and H357-R *POLK* KO cells at different time points after release from double-thymidine block was shown. G1, S, and G2M phases were demarcated (i). The percentages of cells in the G1, S, and G2-M phases at 12 h are estimated and plotted. Data were represented as mean ± SEM (n=2). Statistical significance determined by Ordinary one-way ANOVA, Tukey’s multiple comparisons test, (*p < 0.05, and ns-nonsignificant) (ii).

### DNA polymerase Kappa dependent DNA replication in cisplatin-resistant OSCC cells

Since we observed lower colonization efficiency, reduced proliferation rate, and slower cell-cycle progression in *POLK*-depleted H357-R cells under normal physiological conditions, we hypothesized that DNA replication in these cells might have become *POLK* dependent. To test it further, sensitivity of various OSCC cells to replication inhibitors was determined (**Figure 4A**). While hydroxyurea (HU) depletes the cellular level of dNTPs, aphidicolin inhibits DNA synthesis by binding near the dNTP-binding site of replicative DNA polymerases like Polδ. Camptothecin (CPT) works by inhibiting DNA Topoisomerase I, an enzyme crucial for DNA replication. We found that H357-S and H357-R *POLK* KO exhibited greater lethality to these drugs than H357-R cells, and, interestingly, cells lacking Polκ showed maximal sensitivity, indicating an unexpected role for a TLS Pol during normal DNA replication. Since the processive DNA synthesis during replication in human cells is mostly coordinated by Pols 8 and χ, it is intriguing to determine whether Polκ plays a structural or catalytic role and if its interaction with PCNA becomes important in such an event. In our earlier study, we identified two PIP motifs (528-SIIGFL-533 and 864-TLDIFF-869) in Polκ required for PCNA binding, and showed that TLS occurs normally in human fibroblast cells in which the *pip1* or *pip2* mutant Polκ is expressed, but mutational inactivation of both PIPs rendered Polκ nonfunctional in TLS (Yoon et al. 2014). To delineate the importance of the catalytic domain and PIP motifs of Polκ in DNA replication of OSCC cells, WT, catalytically dead (D198A, E199A), and PIP mutant (F532A, L533A, F868A, F869A) of Polκ were transfected into H357-R *POLK* KO cells, and the susceptibility of the transfectants to replication inhibitors was determined (**Figure 4 B and C**). By probing with an anti-GFP antibody, we first confirmed that all transfectants expressed Polκ at similar levels, except for the vector control (**Figure 4B**). The cytotoxicity assays in the presence of HU, Aphidicolin, and CPT revealed that the overexpression of the wild type and catalytically dead mutants of Polκ could suppress the sensitivity of H357-R *POLK* KO cells, but not by overexpressing the PIP mutant, and its sensitivity remained the same as in cells harboring the vector control (**Figure 4C**). These results suggested that Polκ primarily plays a structural role and efficiently facilitates DNA synthesis by the replicative polymerase; however, its interaction with PCNA is also important. To demonstrate a direct role of Polκ in DNA replication, a DNA fiber assay was conducted as depicted in the ray diagram (**Figure 4D**). In this assay, cells are allowed to incorporate nucleotide analogs into nascent DNA during replication, which can be detected by immunostaining, thereby enabling real-time monitoring of several replication parameters, such as speed, stalling, and origin firing at single-molecule resolution (Quinet et al. 2017; Sahu et al. 2024). CldU incorporation into DNA before HU treatment and ldU incorporation during replication restart were followed, and about 300 DNA fibers were monitored (**Supp. Table-1**). The speed of the restarted forks of H357-R (∼ 0.159 μm/min) was almost double than that of H357-S and H357-R *POLK* KO cells (∼0.08 μm/min) (**Supp. Table-1a**). Although the firing of new origins in these cells was unaffected (∼10%), a higher percentage of stalled fork/fork collisions or conversions was detected in H357-S and H357-R *POLK* KO cells (∼30%). Interestingly, the percentage of fork restarts was relatively higher in H357-R cells (∼70%), suggesting that recovery from HU-induced replication stress is more efficient when cells have abundant Polκ (**Supp. Table-1b**). Next, to determine any physical contacts between Polκ and replicative Pols on PCNA that facilitate efficient replication and cellular proliferation, binding between Polκ with Polδ subunits was checked by immunofluorescence and immunoprecipitation. Human Polδ consists of p125, the catalytic, and p50, p68, and p12 accessory subunits (Khandagale et al. 2019; Sahu et al. 2026). Based on the compatibility of the available antibodies, the nuclear localization of Polκ with the p50 and p12 subunits was examined. We observed a higher population of co-localized nuclear puncta of Polκ with p50, and Polκ with p12 proteins in H357-R than in H357-S cells (**Figure 4E**). Further, these interactions were confirmed by co-immunoprecipitation in two ways (**Figure 4F i and ii**). The cellular extract was precipitated with anti-Polκ or anti-PCNA, and Polδ subunits and other proteins were detected with their respective antibodies. In both approaches, we observed interaction between the Polκ and Polδ subunits, and both are bound to PCNA. Immunoprecipitation with IgG beads did not pull down any proteins. In a similar assay, no co-localization foci or interaction between Polk and the non-replicative polymerase Polδ was observed (**Supp. Fig. 3 A and B**). Altogether, these results suggested that Polκ can participate in normal DNA replication in addition to its role as a TLS polymerase by coordinating with replicative DNA polymerases like Polδ. For such a function, the catalytic domain of Polκ becomes dispensable, whereas its interaction with PCNA is sufficient to drive efficient replication. These data established a key tripartite complex of PCNA-Polκ-Polδ that facilitates efficient replication of OSCC-R cells.

**Figure 4:**
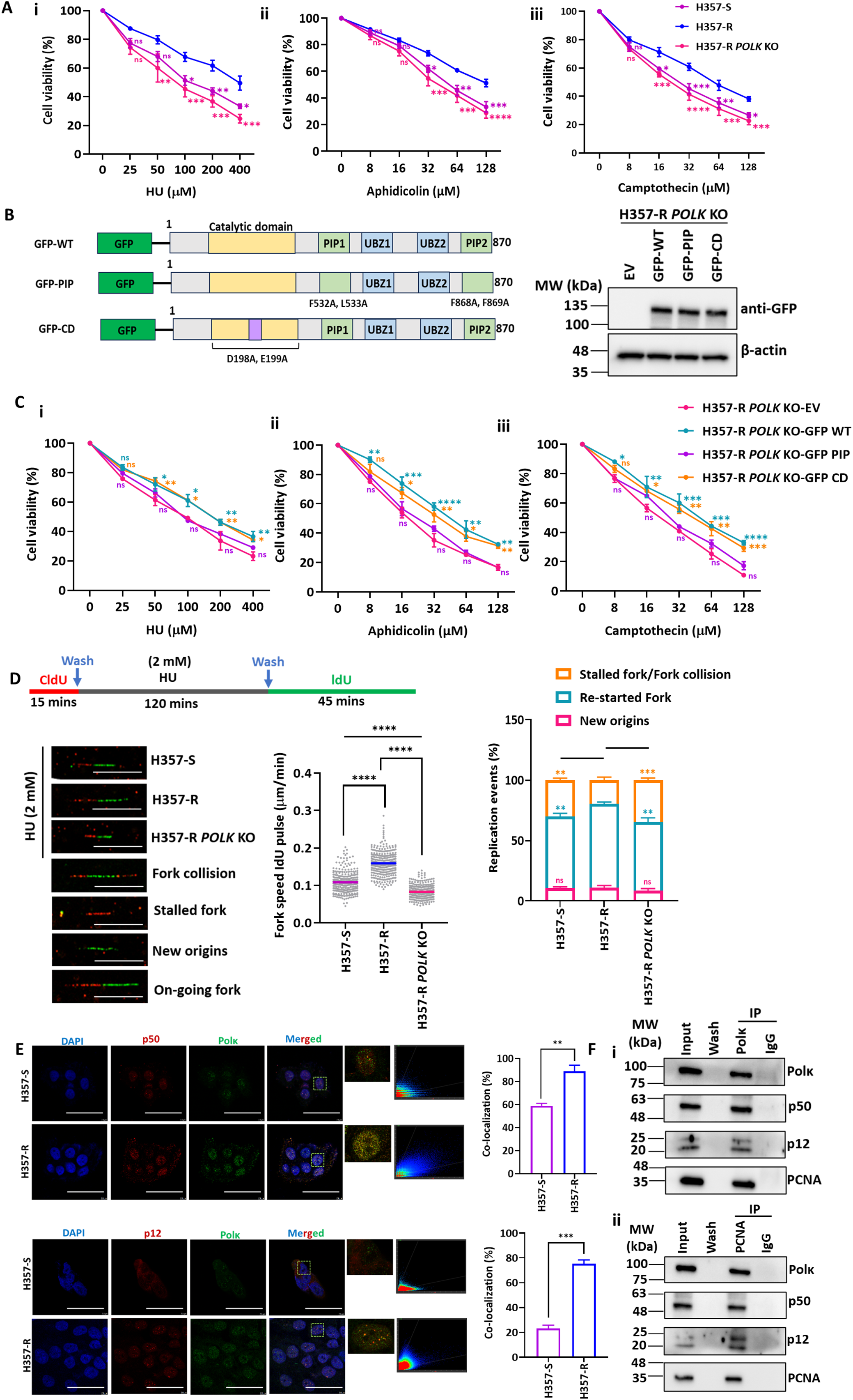
DNA replication in OSCC cisplatin resistant cells is governed by Polκ. **(A)** H357-S, H357-R and H357-R *POLK* KO cells were treated with the indicated concentrations of drugs HU (i), Aphidicolin (ii) and Camptothecin (iii) for 24 hours, and then an MTT assay was performed. These graphs indicate the mean ± SEM (n=3) and were analyzed by two-way ANOVA with Tukey’s multiple comparisons test (*p < 0.05, **p < 0.01, ***p < 0.001, ****p < 0.0001, and ns-nonsignificant). Purple asterisks indicate comparison between H357-R and H357-S; pink asterisks indicate comparison between H357-R and H357-R *POLK* KO. **(B)** A schematic representation of the critical motifs of Polκ and their mutations involved in catalytic activity and PCNA interaction is shown. Knockout cells harbouring GFP-tagged WT, PIP mutant and Catalytically Dead (CD) Polκ were analyzed by Western blotting. EV indicates cells possessing an empty vector. β-actin was taken as an internal control. **(C)** Transfected cells were treated with the indicated concentrations of HU (i), Aphidicolin (ii) and Camptothecin (iii) for 24 hours, and then MTT assays were performed. These graphs indicate the mean ± SEM (n=2) and statistically verified by two-way ANOVA with Tukey’s multiple comparisons test (*p < 0.05, **p < 0.01, ***p < 0.001, ****p < 0.0001, and ns-nonsignificant). Cyan, orange and violet asterisks indicate comparison between EV and GFP-WT, GFP-CD and GFP-PIP, respectively. **(D)** A schematic representation of DNA fibre analysis in the presence of HU and representative images of various replication events in H357-S, H357-R and H357-R *POLK* KO cells were shown. The fork speeds were measured, and the values of three independent experiments were pooled and plotted. Bar graph showing the percentage of various replication events in H357-S, H357-R and H357-R *POLK* KO cells. Data were represented as mean ± SEM (n=3). Statistical significance was determined by two-way ANOVA with Tukey’s multiple comparisons test (**p < 0.01, ***p < 0.001, ****p < 0.0001, and ns-non significant). Scale bars indicate 10 μm. **(E)** Immunofluorescence co-localization analysis of H357-S and H357-R cells showing p12 and p50 subunits of Polδ (red), Polκ (green), DAPI (blue), and corresponding merged images. Merged images represent co-localization of Polδ with Polκ in the nucleus. A zoomed-in merged (only green and red) view of the dashed box area is shown in the upper right corner of the corresponding image. A scatter plot of representative merged images is shown. The percentage of co-localization was shown in a bar diagram. The scatter plot and co-localization percentage were obtained from a confocal microscope Stellaris 5. Data were represented as mean ± SEM (n=3). Statistical significance was determined by t-test, (**p < 0.01, ***p < 0.001). Scale bars indicate 7.5 μm. **(F)** Co-immunoprecipitation was performed in H357-R whole-cell extract using either Polκ or PCNA antibody. Interaction between Polκ and PCNA/p50/p12/Polκ and interaction between PCNA and Polκ/p50/p12/PCNA were analysed by Western blotting. IgG was considered as a negative control. Representative best immunoblots from two independent experiments are shown.

### Catalytic independent function of Polκ is required to protect the replication fork and restart post-cisplatin exposure

Since Polκ depletion sensitized the H357-R cells to cisplatin, we argued that TLS activity by this DNA polymerase could be critical. Although there is no biochemical evidence to suggest nucleotide insertion against cisplatin adducts by Polκ, the catalytic activity may be required during the mismatch extension step in a two-DNA-polymerase-based TLS, where one inserts and Polκ extends, which may be crucial for resistance (Wolfle et al. 2003). To function during TLS, DNA polymerase kappa must also interact with PCNA (Acharya et al. 2020b). To delineate the importance of the catalytic domain and PIP motifs of Polκ in cisplatin resistance of OSCC, after confirming approximately equal expression, the susceptibility of H357-R *POLK* KO cells harboring WT, catalytically dead (Polκ CD: D198A, E199A), and PIP mutant (F532A, L533A, F868A, F869A) of Polκ to cisplatin was determined (**Figure 5A i**). The clonogenic assay confirmed that the wild-type and catalytically inactive mutant Polκ-expressing cells (H357-R, H357-R *POLK* KO + GFP-Polκ, and H357-R *POLK* KO + GFP-Polκ CD) grew well and formed similar numbers of colonies in the presence and absence of cisplatin, again confirming the involvement of Polκ in cisplatin resistance. To our surprise, the cells expressing the PIP mutant of Polκ formed significantly fewer colonies similar to the vector control and this number was further reduced in the presence of cisplatin (**Figure 5A ii and iii**). The MTT assay further confirmed that the ectopic expression of wild-type and catalytically inactive mutants of Polκ reduced sensitivity to cisplatin, and that viability in the presence of cisplatin decreased when cells did not harbor Polκ (EV) or expressed only the PIP mutant (**Figure 5B**). To provide direct evidence of the non-catalytic role of Polκ in cisplatin resistance, GFP-tagged wild-type and mutant forms of Polκ were overexpressed in both H357-S and SCC9-S cells, and cisplatin toxicity was measured by MTT assay (**Supp. Fig. 3C and D**). Notably, cells expressing GFP-Polκ and GFP-Polκ CD exhibited greater tolerance to cisplatin than cells expressing the GFP-Polκ PIP mutant or sensitive cells alone, but showed comparable resistance to H357-R and SCC9-R cells. Since in both cellular model systems the catalytic activity of Polκ was not involved in cisplatin resistance, we conclude that, rather than mutagenic bypass, an alternative structural function of Polκ is critical for exhibiting cisplatin resistance in OSCC cells.

**Figure 5:**
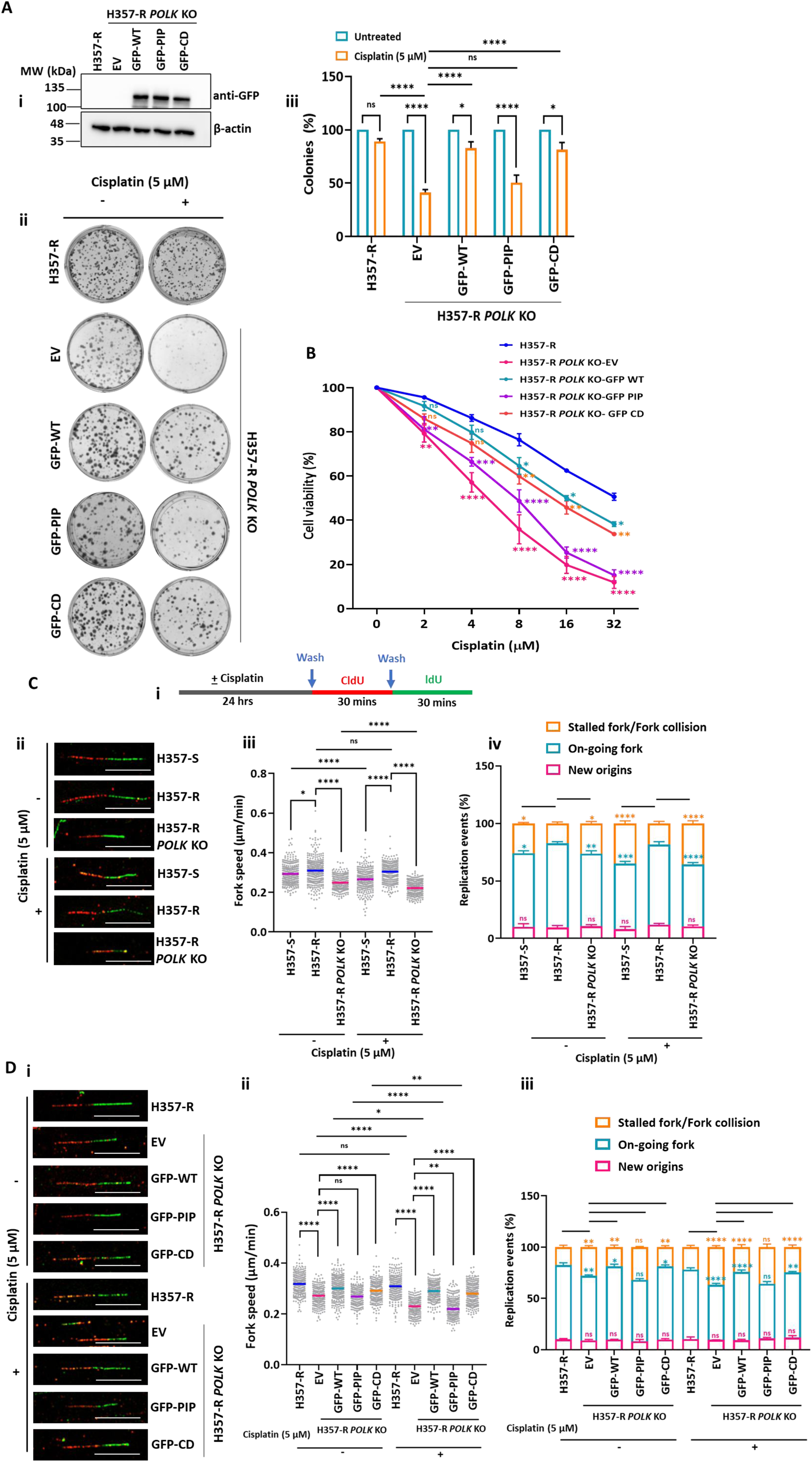
Structural function of Polκ in protecting the replication fork and restart post-cisplatin exposure. **(A)** Western blot analysis shows ectopic expression of GFP-tagged WT, PIP mutant and Catalytically Dead (CD) mutants in H357-R *POLK* KO cells. EV indicates an empty vector. β-actin was considered an internal control (i). A colony-forming assay was performed using H357-R and H357-R *POLK* KO transfected cells (ii). Bar graph represents colony percentage as mean ± SEM (n=4), statistical significance was determined by two-way ANOVA with Tukey’s multiple comparisons test (*p < 0.05, ****p < 0.0001, and ns-nonsignificant) (iii). **(B)** Cells were treated with the indicated concentrations of cisplatin for 48 hours, and then an MTT assay was performed. These graphs indicate the mean ± SEM (n=3), two-way ANOVA, Tukey’s multiple comparisons test (*p < 0.05, **p < 0.01, ***p < 0.001, ****p < 0.0001, and ns-nonsignificant). Pink asterisks indicate comparison between H357-R and H357-R *POLK* KO EV. Cyan, orange and violet asterisks indicate comparison between EV and GFP-WT, GFP-CD and GFP-PIP, respectively. **(C)** Schematic representation of DNA fibre analysis as carried out in the presence and absence of cisplatin (i). Representative replication event images are shown (ii). The fork speeds were measured, and the values of three independent experiments were pooled and plotted (iii). Bar graph showing percentage of various replication events in H357-S, H357-R and H357-R *POLK* KO cells with and without cisplatin (iv). Data were represented as mean ± SEM (n=3). Statistical significance was determined by two-way ANOVA with Tukey’s multiple comparisons test (*p < 0.05, **p < 0.01, ***p < 0.001, ****p < 0.0001, and ns-non significant). Scale bars indicate 10 μm. **(D)** Representative images of DNA fibres in H357-R and H357-R *POLK* KO cells transfected with EV and mutants (GFP-WT, GFP-PIP, GFP-CD) without or with treatment with 5 μM cisplatin for 24 hours before labelling are shown (i). The fork speeds were measured, and the values of three independent experiments were pooled and plotted for H357-R and H357-R *POLK* KO transfected cells with or without cisplatin (ii). The bar graph shows the percentage of various replication events with and without cisplatin. Data are represented as mean ± SEM (n=3) (iii). Statistical significance was determined by two-way ANOVA with Tukey’s multiple comparisons test (*p < 0.05, **p < 0.01, ****p < 0.0001, and ns-non significant). Scale bars indicate 10 μm.

To demonstrate Polκ’s role in replication fork protection and restart upon cisplatin exposure, DNA fiber analysis was carried out. Replication was first blocked with cisplatin, and after washings, cells were allowed to replicate; the DNA fibers were sequentially labelled with two thymidine analogues, detected by immunostaining, and analyzed (**Figure 5C, Supp. Table -2 a and b**). While the speed of the restarted forks of H357-R cells did not change significantly without and with cisplatin treatment (∼0.3 μm/min), in the case of H357-S and H357-R *POLK* KO cells, the fork speeds were much lower (∼0.25-0.29 μm/min) and further reduced under cisplatin stress (∼0.26-0.22 μm/min). Consequently, the percentage of stalled fork/fork collisions (35%) was higher in H357-S and H357-R *POLK* KO cells than that in H357-R cells (∼15-24%). Evidently, more actively replicating forks (∼66-71%) were found in H357-R cells. The replication defects in the absence and presence of cisplatin in H357-R *POLK* KO cells were rescued by overexpressing catalytically active as well as inactive mutants of Polκ (H357-R *POLK* KO + GFP-Polκ, and H357-R *POLK* KO + GFP-Polκ D198A, E199A), but not by the PIP mutant (H357-R *POLK* KO + GFP-Polκ pip) (**Figure 5D and Supp. Table -3**). These results suggested Polκ induces cisplatin resistance in H357 in a catalytically independent manner, but its interaction with PCNA is critical in protecting the replication fork and recovering thereafter. Moreover, the DNA fiber assay without cisplatin further confirmed that the replication defects in H357-R *POLK* KO cells are solely due to the loss of Polκ, which can be rescued by a catalytically dead mutant of Polκ. Thus, the non-TLS function of Polκ plays an important role in replicating both undamaged and cisplatin adducts containing the genome, thereby maintaining uninterrupted DNA replication in chemoresistant OSCC.

### Polκ depletion impairs ATM-ATR signaling and DNA strand-break repair pathways

To delineate the underlying mechanisms by which Polκ non-catalytically protects the genome from cisplatin damage, we were intrigued to explore various DDR pathways beyond TLS. The ATM and ATR pathways are central to the DNA damage response (DDR), which protects genomic integrity by sensing DNA damage, coordinating cell cycle checkpoints, and activating DNA repair mechanisms. While ATM is primarily activated by DSBs to activate downstream kinase Chk2, ATR responds to a broader range of DNA damage, including stalled replication forks, to activate Chk1 to execute the appropriate DNA repair pathways for cell survival, and failing to do so can lead to apoptosis (Casper et al. 2002; Hirao et al. 2002). We found that the protein levels of ATM, ATR, and their phosphorylated forms were remained unaltered in all of these cells H357-S, H357-R and H357-R *POLK* KO without and with cisplatin treatment (**Figure 6A and Supp. Fig. 4A**). However, barely any Chk1 and Chk2 proteins were expressed in Polκ depleted cells, accordingly no activated forms of these proteins (p-Chk1 and p-Chk2) were detected upon cisplatin damage (**Figure 6A and Supp. Fig. 4A**). Whereas a higher expression of p-Chk1 and p-Chk2 were detected in H357-S and H357-R with cisplatin treatment in comparison to untreated cells. This result indicated that activated ATM-ATR kinases can transduce the DDR response in Polκ-proficient cells but are inefficient at doing so in Polκ-deficient cells. ATM-Chk2 regulates DSB repair by homologous recombination (HR) and non-homologous end joining (NHEJ) pathways (Kumari et al. 2025). Mre11-Rad50-Nbs1 (MRN) complex, upon binding to broken DNA, recruits and activates the ATM-Chk2 pathway. Further, they recruit CtIP to DSBs and phosphorylate it to facilitate resection. Chk2 activation leads to BRCA1 phosphorylation, aiding in the recruitment of Rad51 to DNA double-strand breaks. This action facilitates the proper assembly of the Rad51-BRCA2-ssDNA complex to execute the error-free HR repair pathway. RAD54 helps the Rad51-coated DNA search for a homologous template sequence, a critical step for strand invasion. While HR is mostly an error-free repair pathway that occurs in S phase, NHEJ is an intrinsically error-prone repair pathway that occurs at all cell cycle stages. ATM phosphorylates core NHEJ factors, including DNA-PKCs, Artemis, XRCC4, and XLF, as well as DNA polymerase λ (Polλ), thereby promoting NHEJ. We assessed the status of HR and NHEJ proteins in these cells (**Figure 6 B and C**). We observed that while the MRN complex seems to be operational, the downstream proteins of HR are mostly missing only in H357-R *POLK* KO cells without and with cisplatin treatment (**Figure 6 B and Supp. Fig. 4B**). Similarly, NHEJ pathway is also affected as some of the key downstream effectors like phospho-DNA-PKCs, Artemis, and DNA ligase IV were hardly detected in cisplatin treated H357-R *POLK* KO cells (**Figure 6 C and Supp. Fig. 4C**). Again, the upstream factors of NHEJ pathway like Ku70-Ku80 heterodimer complex seems to be active in these cells. We reconfirmed these observations by taking nuclear fractions and Western analyses with specific antibodies, and histone H3 as a control (**Figure 6D and Supp. Fig. 4D**). Nuclear localization analyses by immunostaining further strengthened our observations (**Supp. Fig. 5**). Altogether, these results suggested that Polκ somehow regulate stabilization of downstream factors of cell cycle checkpoints, HR and NHEJ repair pathways to stabilize the replication fork and the genome.

**Figure 6:**
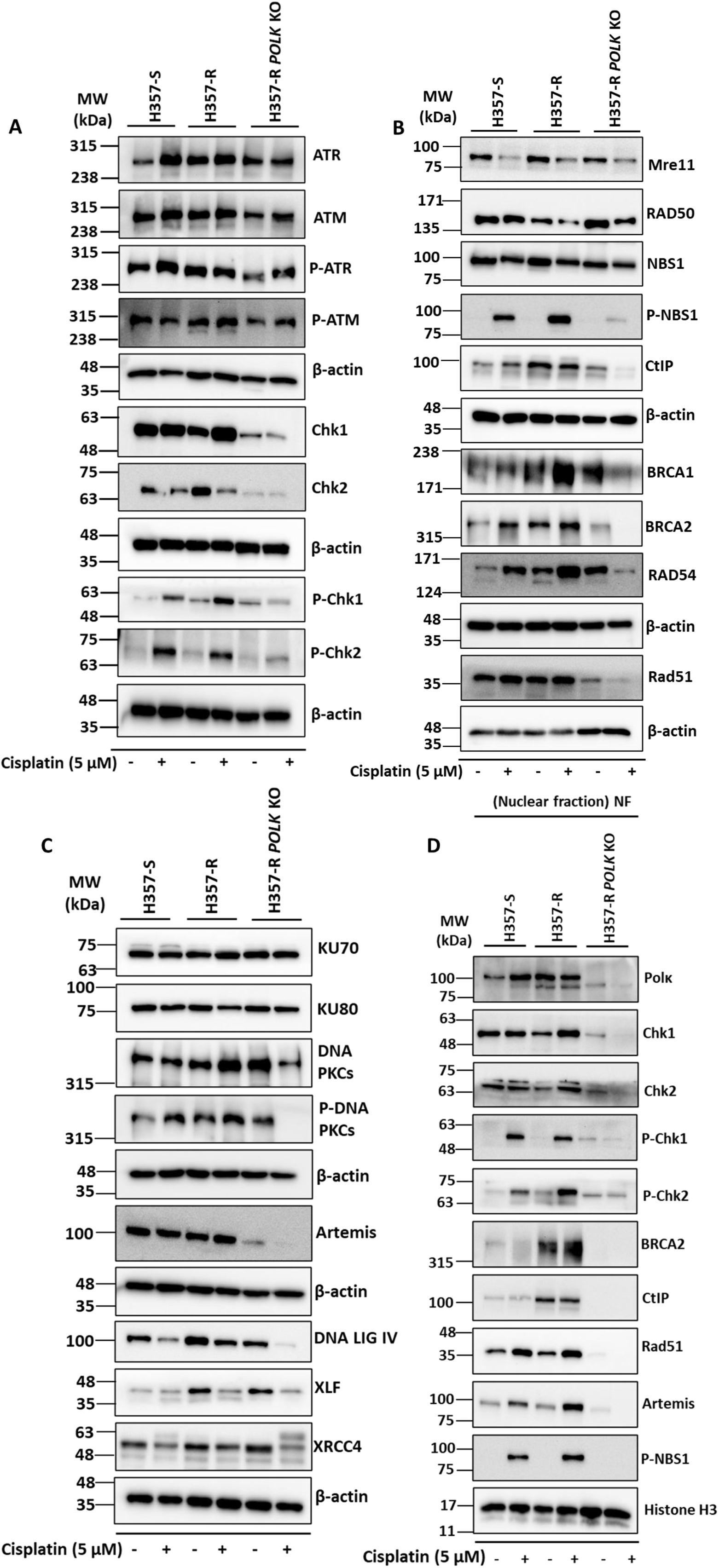
Absence of Polκ dysregulate ATM -ATR signaling and DNA strand break repair pathways. **(A)** Western blot analyses of cell-free lysates of H357-S, H357-R and H357-R *POLK* KO cells without and treated with 5 μM cisplatin were carried out and were probed with the indicated antibodies against checkpoint proteins. β-actin was detected as a loading control. **(B)** Similar Western blot analyses were performed, but probed with antibodies against proteins involved in the HR pathway. **(C)** Similar Western blot analyses were performed, but probed with antibodies against proteins involved in the NHEJ pathway. **(D)** Western blot analyses of nuclear extracts of H357-S, H357-R and H357-R *POLK* KO cell lines without and treated with 5 μM cisplatin were carried out and were probed with the indicated specific antibodies. Histone H3 was used as a loading control.

### Polκ dependent conserved pathways are involved in cisplatin resistance of SCC9 cells

To further strengthen our observations, we used another OSCC cell line, SCC9, to demonstrate the role of Polκ in cisplatin resistance. Since we could not select *POLK^-/-^* clonal cells by using the CRISPR-Cas9 approach, we decided to use the siRNA approach to knock down *POLK* and characterize such cells further (**Supp. Fig. 6**). First, we confirmed the efficacy of *POLK* siRNA, and it is able to knock down most of the transcripts, as we hardly detected any Polκ protein by Western and immunofluorescence analyses (**Supp. Fig. 6A i and Supp. Fig. 6B**). Cell viability assay suggested that the depletion of Polκ re-sensitizes SCC9-R resistant cells to cisplatin and the IC50 value reduced from ∼32 µM to ∼8 µM of cisplatin (**Supp. Fig. 6D**). Comet assay, ψ-H2A.X, and cleaved PARP1 detection further ascertained the role of Polκ in protecting genome of SCC9 from damages caused by cisplatin (**Supp. Fig. 6 C, E, and Supp. Fig. 6 A ii and iii**). The absence of Chk1, Chk2, p-Chk1, p-Chk2, CtIP, BRCA2, Rad51, and Artemis in *POLK* knockdown SCC9 cells, with and without cisplatin treatment, suggested a possible critical role of Polκ in activating and transducing cell-cycle checkpoints to regulate HR and NHEJ to stabilize the genome (**Supp. Fig. 6A iv-xi**). These results confirmed that Polκ plays conserved roles in inducing cisplatin chemoresistance, likely also in all OSCC cell types.

### Polκ protects key checkpoint and DSB repair proteins from ubiquitin-mediated degradation

Low-level detection of checkpoint and DDR proteins in Polκ-depleted cells could be due to defects at the gene expression or post-translational levels. Therefore, RNA expression in terms of cDNA was checked by agarose gel electrophoresis (**Figure 7A i and ii**). We observed cDNAs of transcripts encoded by *CHEK1*, *CHEK2*, *BRCA2*, *CTIP*, *RAD51*, and *ARTEMIS* genes in both Polκ proficient and deficient H357-R cells, whereas, as expected, no cDNA was detected for *POLK* in H357-R *POLK*^-/-^ cells. Next, we assessed the expression of these proteins in the presence of MG132, an inhibitor that blocks the proteolytic activity of the 26S proteasome, which degrades unneeded or damaged proteins. Our analyses suggested that MG132 has no effect on the protein levels in H357-R cells, whereas Chk1, Chk2, BRCA2, CtIP, Rad51, and Artemis proteins were recovered in H357-R *POLK* KO cells upon MG132 treatment, and barely any of these proteins were detected without the proteasomal inhibitor (**Figure 7B**). The results were further consolidated by immunofluorescence, as the nuclear localization of Chk1, Chk2, BRCA2, Rad51, and Artemis increased in MG132 treated H357-R *POLK* KO cells (**Figure 7C and Supp. Fig. 7A**). Thus, the absence of protein bands in Polκ depleted cells is most likely due to proteasomal degradation, and Polκ could enhance the stability and cellular availability of these critical DDR proteins.

**Figure 7:**
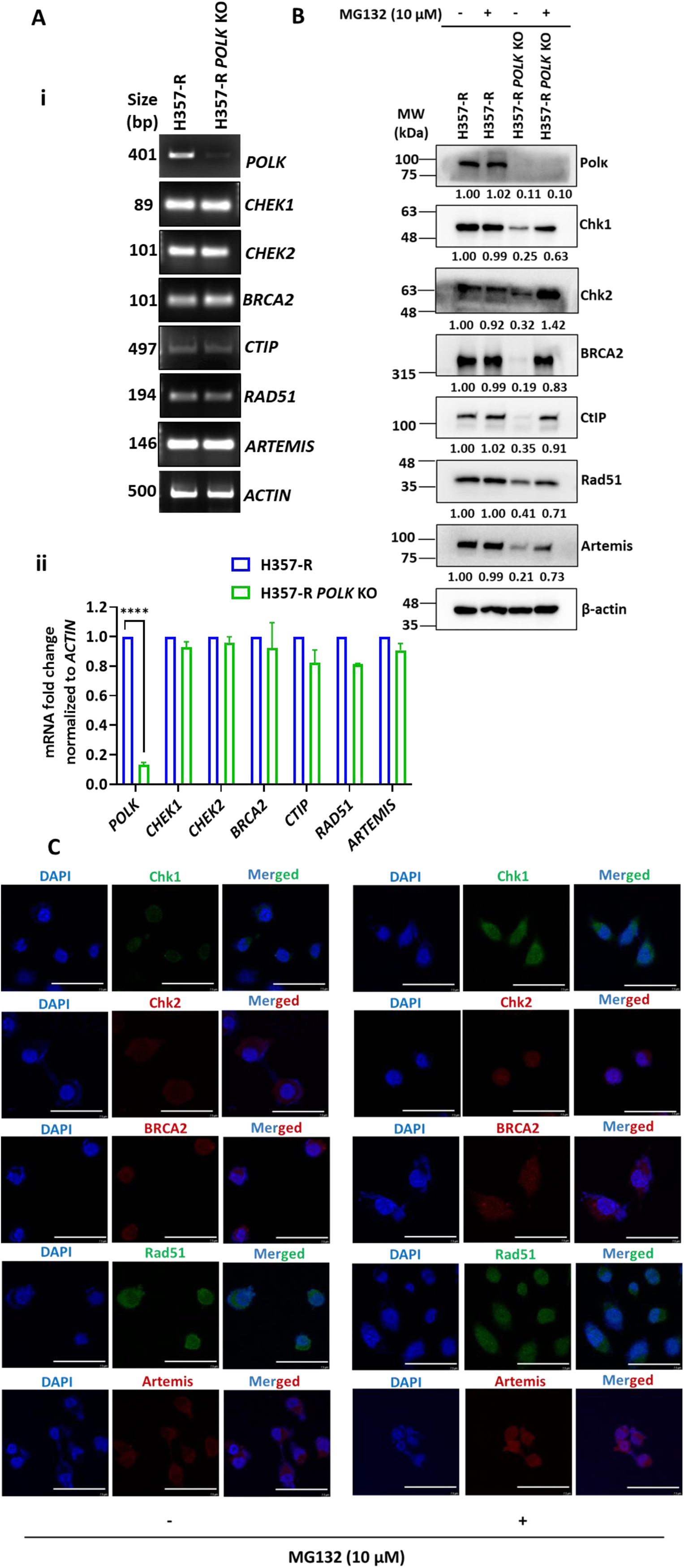
Stabilization of proteins involved in checkpoint and DSB repair pathways from proteasomal degradation by Polκ. **(A)** RT-PCR analysis of indicated genes was carried out in H357-R and H357-R *POLK* KO cells (i), and the mRNA expression levels were represented in a bar graph (ii). Data were represented as mean ± SEM (n=2). Statistical significance determined by two-way ANOVA with Tukey’s multiple comparisons test (****p < 0.0001, and ns-nonsignificant). **(B)** Western blot analyses of cell lysates of H357-R *POLK* KO and H357-R *POLK* KO cells without and treated with 10 μM MG132 for 6 hours were conducted and probed with the indicated antibodies. β-actin was used as a loading control. Densitometric fold change relative to proteins in H357-R cells has been provided just below each band. **(C)** Representative immunofluorescence images of H357-R *POLK* KO and H357-R *POLK* KO cells treated with 10 μM of MG132 show DAPI (blue), Chk1 (green), Rad51 (green), Chk2 (red), BRCA2 (red), and Artemis (red), and corresponding merged images. Scale bars indicate 7.5 μm.

### Dysregulation of proteasomal degradation and deubiquitinating systems in cisplatin resistant OSCC

To delineate the possible molecular mechanism by which Polκ protects against ubiquitin-mediated degradation of critical DDR proteins, we reanalyzed publicly available RNA-seq data from H357-S and H357-R cells [ArrayExpress (E-MTAB-9697)]. The total number dysregulated genes were accessed, and STRING analysis was carried out. Interestingly, in addition to several altered pathways, around 349 genes involved in the protein degradation pathway were dysregulated (**Figure 8A i-iv, Supp. Table-4 a and b**). Of these, 76 and 8 genes involved in ubiquitination pathways (e.g., *UBC, ISG15, RNF185*, and *ATG10*, etc.) and 20S proteasomal complex formation (*PSMB5, PSMB10, PSMC1, PSMC5, PSMD6, PSMD9, PSMD12,* and *PSME3*) were significantly down-regulated in H357-R compared with H357-S cells. Meanwhile, 13 genes, including *USP2, USP18, USP42, USP43*, and *OTUD3*, involved in the removal of ubiquitin from its conjugates were up-regulated in H357-R cells. The remaining genes were marginally dysregulated. Our RNA-seq analyses indicated that protein degradation pathways are likely affected in cisplatin resistant cells due to overall downregulation of the 20S proteasomal complex and upregulation of deubiquitinating enzymes. To validate this, semi-quantitative and real-time RT-PCRs were performed for two deubiquitinase genes, *USP2* and *USP18*, and two proteasomal complex genes *PSMA7* and *PSMB5* (**Figure 8B i and ii)**. While *USP2* encodes a ubiquitin-specific protease of the C19 peptidase superfamily which deubiquitinates polyubiquitinated target proteins such as fatty acid synthase, MDM2, MDM4, and cyclin D1 (Davis et al. 2016; Tang et al. 2022; Zhu et al. 2022; Li et al. 2026), *USP18* (alias *UBP43*) encodes a major ISG15-specific protease that efficiently and selectively cleaves ISG15 (a ubiquitin-like protein) fusions (D’Andrea and Pellman 1998; Geng et al. 2024). USP18 regulates several biological processes, including immune cell development, autoimmune diseases, pathogen responses, and tumor development (Honke et al. 2016). *PSMB5* (Proteasome Subunit Beta Type-5) encodes a catalytic subunit of the 20S proteasome core complex, crucial for protein degradation in eukaryotes (Oerlemans et al. 2008). *PSMA7* (Proteasome Subunit Alpha Type-7) encodes a critical structural component of the 20S proteasome core complex and, as an alpha subunit, forms the “gateway” of the proteasome, regulating the entry of proteins into the catalytic core for degradation (Wu et al. 2021). We selected these four genes for further analysis because they are highly altered and reported to promote the development, progression, and chemoresistance of several tumours. Total cDNA was synthesised from the RNA isolated from H357-S, H357-R, and H357-R POLK KO cells. Endpoint reverse transcription and qRT-PCR results revealed ∼2-4-fold higher expression of USP2 and USP18 in H357-R cells. *PSMB5* and *PSMA7* mRNA expression was somewhat lower in H357-R cells but ∼3-fold higher in H357-*POLK* KO cells. To further strengthen this, the expression of these proteins was assessed using specific antibodies (**Figure 8C**). H357-R cells exhibited increased expression of USP2 and USP18 and reduced levels of PSMA7 and PSMB5 compared to other cells. These findings suggest that decreased ubiquitination, together with enhanced USP2/USP18 deubiquitinase activity, may contribute to the stabilization of key cell cycle checkpoint and DNA damage response (DDR) proteins. This, in turn, likely facilitates replication fork protection and restart, thereby promoting genome stabilization and cisplatin resistance in OSCC-R cells.

**Figure 8:**
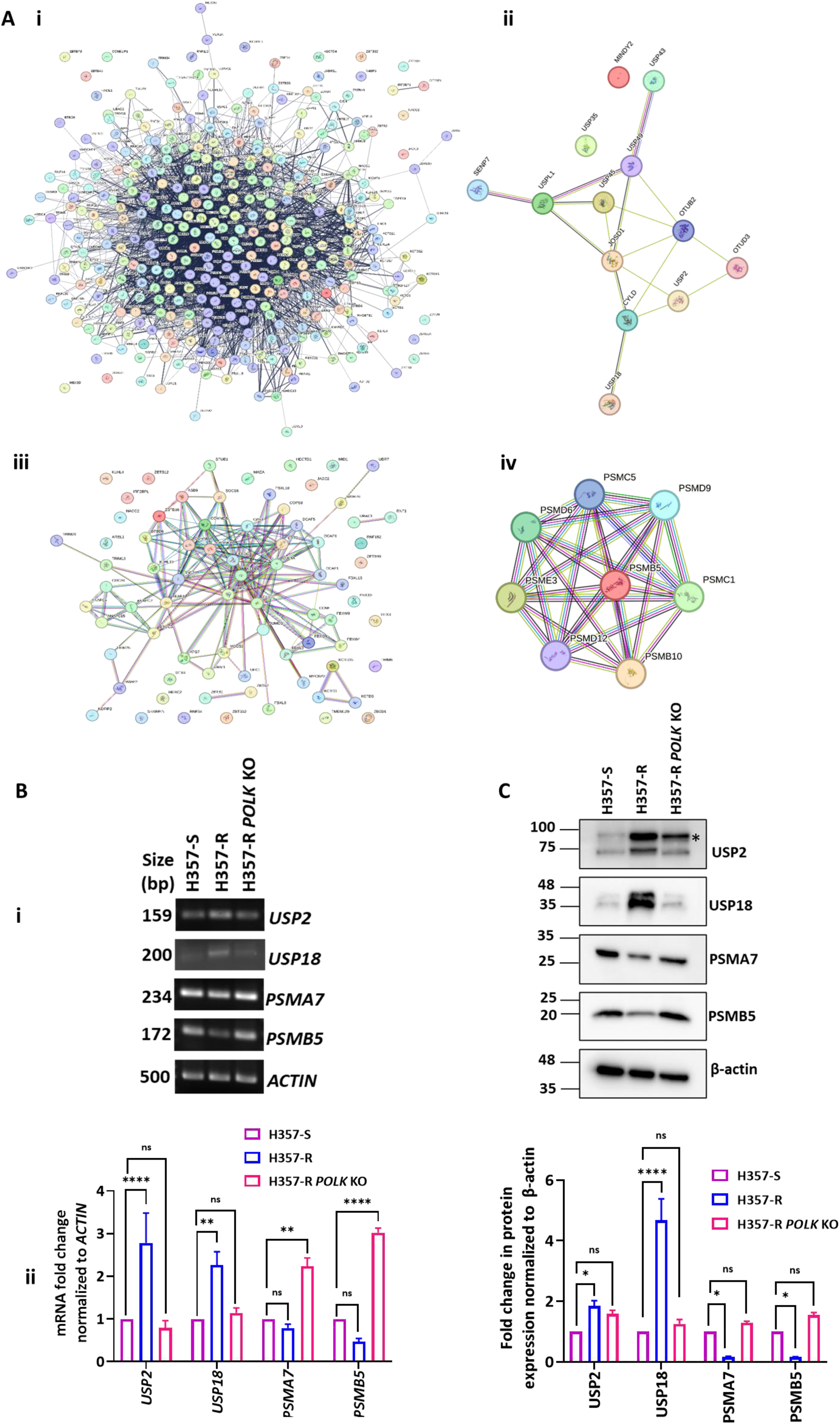
Dysregulation of proteasomal degradation and deubiquitinating systems in cisplatin resistant OSCC. **(A)** Comparative transcriptomic data of H357-S and H357-R were retrieved and re-analysed. String analysis revealed a total of 349 dysregulated genes in protein degradation pathways (i), out of which 13 genes are significantly up-regulated in deubiquitination pathways (ii), 76 genes are down-regulated and involved in ubiquitination pathways (iii), and 8 genes are down-regulated; these are components of the 26S proteasomal complex (iv). **(B)** Endpoint RT-PCR analysis of the indicated genes was carried out in H357-S, H357-R and H357-R *POLK* KO cells (i) and qRT–PCR gene expression analysis of the indicated genes in H357-S, H357-R and H357-R *POLK* KO cells was performed using SYBR Green. ACTIN was used as a control. ΔΔct values were calculated and compared for the H357-S, H357-R and H357-R *POLK* KO cells. Calculated fold change was plotted. Data were represented as mean ± SEM (n=3) (ii). Statistical significance was determined by two-way ANOVA with Tukey’s multiple comparisons test (**p < 0.01, ***p < 0.0001, and ns-nonsignificant). **(C)** Western blot analyses of H357-S, H357-R and H357-R *POLK* KO lysate were carried out and probed with the indicated antibodies. β-actin was used as a loading control. (*) indicates a non-specific band for the USP2 antibody. The USP18 antibody detected two protein sizes. Fold change in protein expression was shown in a bar graph. Data are represented as mean ± SEM (n=2). Statistical significance was determined by two-way ANOVA with Tukey’s multiple comparisons test (*p < 0.05, ****p < 0.0001, and ns-nonsignificant).

### PCNA-Polκ-USP18 axis plays an important role in chemoresistance in OSCC

Next, we assessed for any physical and functional interactions between Polκ and deubiquitin enzymes that regulate the stabilization of critical DDR proteins and cisplatin resistance in OSCC using immunoprecipitation and nuclear co-localization. The complete cellular extract was either precipitated with anti-Polκ or anti-PCNA antibodies, and the USP2 and USP18 proteins were detected with their respective antibodies. While immunoprecipitation with IgG beads did not precipitate any proteins, in both approaches, we could observe a positive interaction between Polκ, PCNA, and USP18 (**Figure 9A and Supp. Fig. 7B**). Immunoprecipitation either with USP2 or Polκ antibodies could not pull down the other partner, suggesting no interaction between the two. Moreover, we could not detect an interaction between USP2 and PCNA. The nuclear co-localization analysis also confirmed similar results (**Figure 9B**). While the binding between PCNA and Polκ is well known to be mediated by their hydrophobic interdomain connecting loop (IDCL) and PIP motif, respectively, the binding of USP18 to PCNA or Polκ may be indirect. Polκ possesses two ubiquitin binding Zn finger motifs (UBZ1 and UBZ2) juxtaposed between two PIP motifs, and such motifs are known to interact with proteins [**Fig. 9C i** (Bienko et al. 2005; Manohar et al. 2022)]. To verify the mode of interaction among these three proteins, cell lysates expressing GFP-Polκ, GFP-Polκ PIP, and GFP-Polκ UBZ were immunoprecipitated with a USP18 antibody, and the partner proteins were detected using anti-GFP and anti-PCNA antibodies (**Fig. 9C ii-iv)**. Interestingly, while USP18 could pull down both wild type Polκ and PCNA, it failed to precipitate either protein from cells expressing Polκ with PIP or UBZ mutations. To further strengthen this, we used an anti-PCNA antibody to pull down GFP-UBZ and USP18. Immunoprecipitation results suggested that PCNA could interact with the UBZ mutant of Polκ but failed to precipitate USP18 (**Fig. 9C v**). This result suggested that while PCNA binds to the PIP motifs, USP18 binds to the UBZ domains of Polκ. It also suggested that USP18 interacts only with the Polκ-PCNA complex, and that binary complexes of Polκ-USP18 and PCNA-USP18 are unlikely to exist. Since the UBZ of Polκ appear to be as critical as the PIP motif for the PCNA-Polκ-USP18 axis, we wanted to check its functional relevance in the context of cisplatin resistance and replication continuity. MTT assay confirmed that overexpression of the UBZ mutant of GFP-Polκ could not suppress the sensitivity of H357-R POLK^-/-^cells (**Fig. 9 D i and ii**). DNA fibre analyses also confirmed that replication fork speed is affected in the cells expressing the UBZ mutant of Polκ, which is just as good as H357-R POLK^-/-^ cells, and accordingly, a higher percentage of stalled and fork conversions fibrer were detected (**Fig. 9 E i - iii).** Altogether, we demonstrated the existence of a trimeric axis of PCNA-Polκ-USP18, in which USP18 is recruited to the PCNA bound Polκ, and the both the PIP and UBZ motifs of Polκ are indispensable for cisplatin resistance and replication fork rescue, especially under cisplatin induced genotoxic stress.

**Figure 9:**
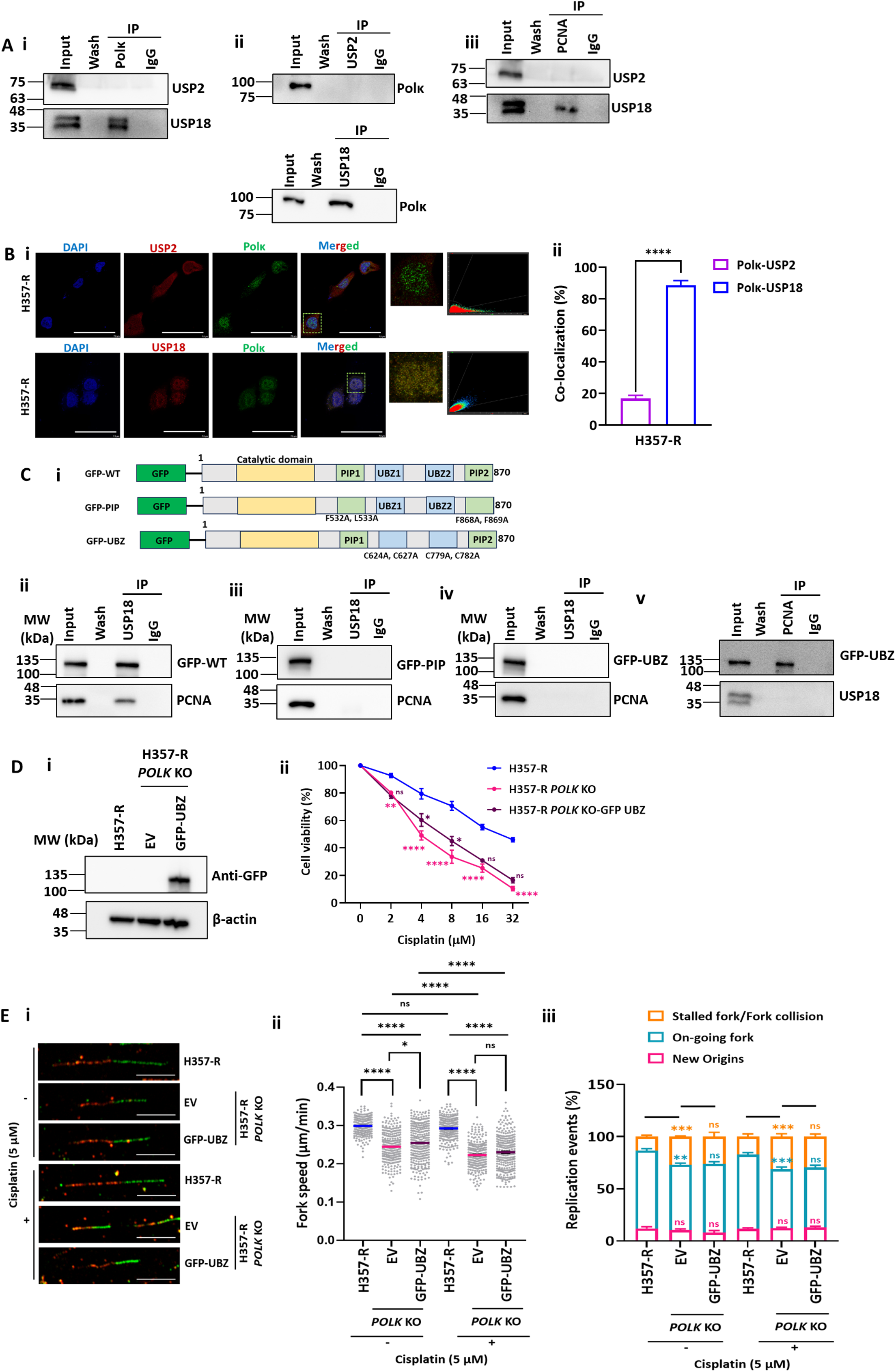
PCNA-Polκ-USP18 axis is critical for cisplatin resistance. **(A)** Co-immunoprecipitation was performed in H357-R whole-cell extract using either Polκ (i) or USP2 and USP18 antibodies (ii). Similar interactions between PCNA and USP2 and USP18 were determined (iii). IgG was considered as a negative control. Representative best immunoblots from two independent experiments are shown. **(B)** Immunofluorescence co-localization analysis of H357-R cells showed USP2 (red), USP18 (red), Polκ (green), DAPI (blue), and corresponding merged images. Merged images represent co-localization of USP2 and USP18 with Polκ in the nucleus. A zoomed-in merged (only green and red) view of the dashed box area is shown in the upper right corner of the corresponding image. A scatter plot of representative merged images is shown (i). The percentage of co-localisation was shown in a bar diagram. The scatter plot and co-localization percentage were obtained from a confocal microscope, Stellaris 5. Data were represented as mean ± SEM (n=3). Statistical significance was determined by t-test, (****p < 0.0001). Scale bars indicate 7.5 μm (ii). **(C**) A schematic representation of Polκ depicting PIP and UBZ motifs and their mutations is shown (i). H357-R *POLK* KO cells were transfected with GFP-WT, GFP-PIP, and GFP-UBZ constructs. After 48 hours of transfection, cells were collected, and Co-immunoprecipitation was performed in GFP-WT (ii), PIP (iii), and UBZ (iv) overexpressing cell extracts using USP18 antibody and pull-down of Polκ and PCNA was verified. Using a similar extract, anti-PCNA antibody was used to precipitate Polκ UBZ mutant and USP18 (v). IgG was considered as a negative control. Representative immunoblots from two independent experiments are shown. **(D)** Western blot analysis shows ectopic expression of GFP-tagged UBZ mutant in H357-R *POLK* KO cells. EV indicates an empty vector. β-actin was used as an internal control (i). Cells were treated with the indicated concentrations of cisplatin for 48 hours, and then an MTT assay was performed. These graphs indicate the mean ± SEM (n=3), two-way ANOVA, Tukey’s multiple comparisons test (*p < 0.05, **p < 0.01, ****p < 0.0001, and ns-nonsignificant). Pink asterisks indicate comparison between H357-R and H357-R POLK *KO* with EV. Burgundy asterisks indicate comparison between EV and GFP-UBZ (ii). **(E)** Representative images of DNA fibres in H357-R and H357-R *POLK* KO cells transfected with EV and GFP-UBZ mutant without or with treatment with 5 μM cisplatin for 24 hours before labelling are shown (i). The fork speeds were measured, and the values from three independent experiments were pooled and plotted for H357-R and H357-R *POLK* KO-transfected cells with or without cisplatin (ii). The bar graph shows the percentage of various replication events with and without cisplatin. Data are represented as mean ± SEM (n=3) (iii). Statistical significance was determined by two-way ANOVA with Tukey’s multiple comparisons test (**p < 0.01, ***p < 0.001, and ns-non significant). Scale bars indicate 10 μm.

### USP18 of the trimeric axis plays an important role in DDR proteins stabilization and chemoresistance in OSCC

To investigate how USP18 depletion could affect protein stability, USP18 expression was silenced in H357-R cells using specific siRNA, followed by analysis of key DNA damage response (DDR) proteins by Western blotting. We observed that upon knocking down *USP18,* the protein stability of Chk1, Chk2, CtIP, and Artemis decreased significantly, similar to Polκ depletion, but recovered only when a proteasomal inhibitor was added to the cells (**Figure 10 A and B**). This result suggested that loss of *POLK* or *USP18* in H357-R cells leads to renders to a similar fate for DDR proteins, likely affecting replication fork protection and restart in the presence of cisplatin. MTT assays further showed that cisplatin sensitivity was enhanced upon *USP18* knockdown (**Figure 10 C**). DNA fibre analyses also confirmed that replication fork speed is affected in the cells upon *USP18* knockdown, as in H357-R POLK^-/-^ cells, and accordingly, a higher percentage of stalled and fork conversions fibres was detected, and the effect was more pronounced in the cisplatin-treated cells (**Fig. 10 D i - iii).**

**Figure 10:**
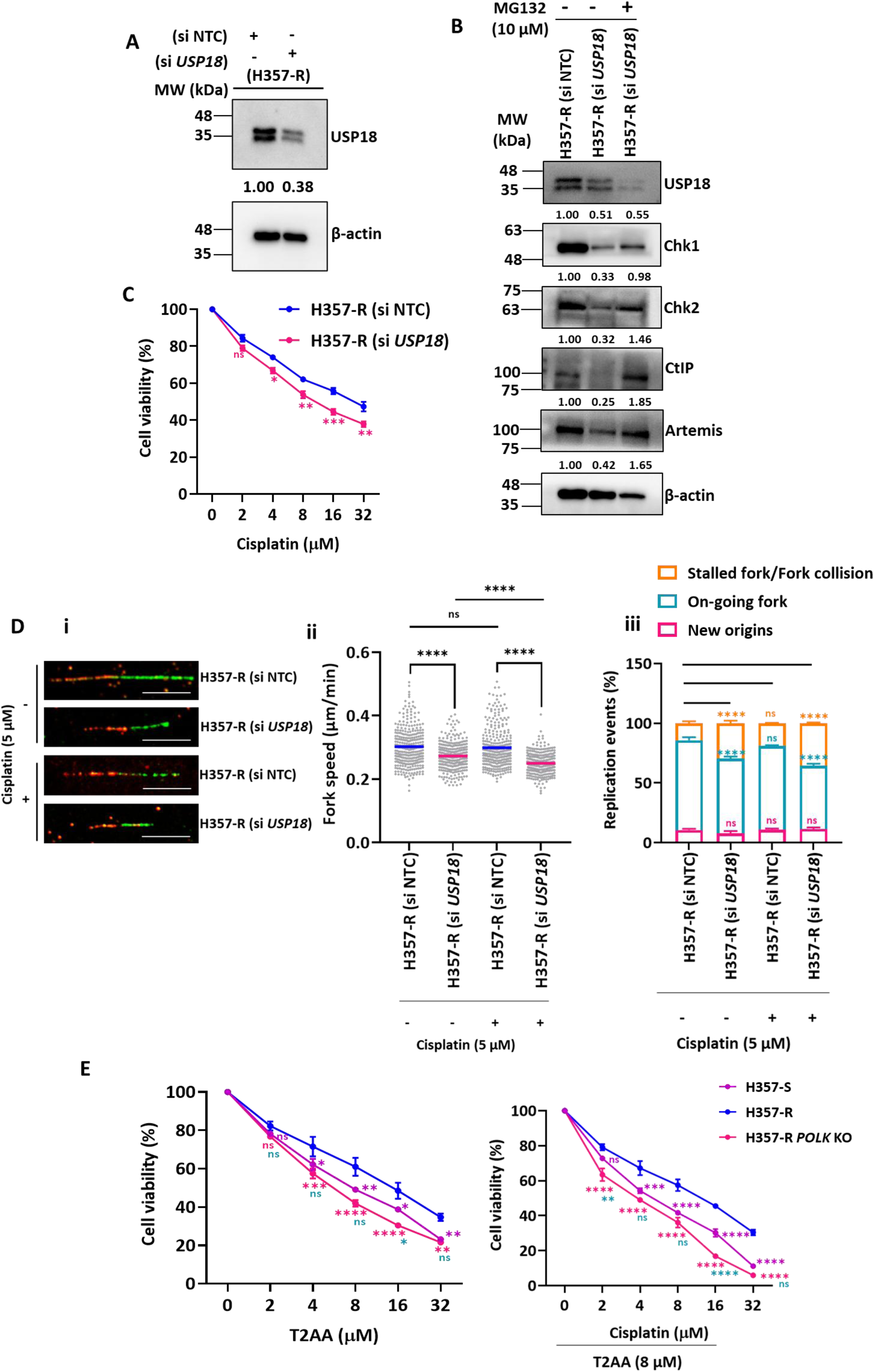
PCNA-Polκ-USP18 axis is critical for cisplatin resistance. **(A)** H357-R cell lines were transfected with NTC siRNA and *USP18* siRNA as described in the Methods. Lysates were collected from the indicated cells, and Western blotting was performed with the indicated antibodies. β-actin was used as an internal control. Representative best immunoblots from two independent experiments are shown. Densitometric fold change relative to proteins in H357-R cells has been provided just below each band. **(B)** Western blot analyses of cell lysates of H357-R, MG132-untreated, and treated H357-R *USP18* knockdown cells were carried out, and expression levels of indicated proteins were determined. Densitometric fold change relative to β-actin has been provided just below each band. **(C)** Cells were treated with the indicated concentrations of cisplatin for 48 hours, and then an MTT assay was performed. The graph indicates the mean ± SEM (n=3), and statistical significance was determined by two-way ANOVA, Šídák’s multiple comparisons test, (*p < 0.05, **p < 0.01, ***p < 0.001, and ns-nonsignificant). Pink asterisks indicate comparison between H357-R (si NTC) and H357-R (siUSP18). **(D)** Representative images of DNA fibres in H357-R (siNTC) and H357-R (siUSP18) cells without or with treatment with 5 μM cisplatin for 24 hours before labelling are shown (i). The fork speeds were measured, and the values from three independent experiments were pooled and plotted for H357-R (si NTC) and H357-R (siUSP18) cells, with or without cisplatin (ii). The bar graph shows the percentage of various replication events with and without cisplatin. Data were represented as mean ± SEM (n=3) (iii). Statistical significance was determined by two-way ANOVA with Tukey’s multiple comparisons test (****p < 0.0001, and ns-non significant). Scale bars indicate 10 μm. **(E)** H357-S, H357-R and H357-R *POLK* KO cells were treated with the indicated concentrations of T2AA alone and with cisplatin for 48 hours, and then an MTT assay was performed. Graph indicates the mean ± SEM (n=3), two-way ANOVA with Tukey’s multiple comparisons test (*p < 0.05, **p < 0.01, ***p < 0.001, ****p < 0.0001, and ns-nonsignificant). Purple asterisks indicate comparison between H357-R and H357-S, pink asterisks indicate comparison between H357-R and H357-R *POLK* KO, and green asterisks indicate comparison between H357-S and H357-R *POLK* KO.

Since Polκ appears to be the master regulator of several pathways that induce chemoresistance, targeting this TLS Pol could improve the efficacy of cisplatin. As Polκ’s interaction with PCNA is indispensable for such a role, the binding site between the two proteins may be targeted. We leveraged an existing small molecule T2AA that binds to the interdomain connecting loop (IDCL), where most DNA Pols typically bind (Punchihewa et al. 2012; Inoue et al. 2014). Polκ binds to the IDCL of PCNA; since T2AA is a generic inhibitor and can prevent binding of all other Pols, it inhibited cell survival of H357-S, H357-R, and H357-R *POLK* KO cells, *albeit* more in H357-S and H357-R *POLK* KO cells (**Figure 10 E**).

The addition of cisplatin further heightened the sensitivity of H357-S and H357-R *POLK* KO cells. A specific small molecule that can disrupt PCNA-Polκ interaction will be much more relevant for clinical use. Also, an inhibitor of USP18 may enhance the reversal of cisplatin resistance in OSCC cells, a finding that warrants further small molecules screening and detailed investigation. Nevertheless, this study established yet another tripartite axis of PCNA-Polκ-USP18 that is involved in stabilizing the genome of a highly proliferating cell under cisplatin stress and can be targeted to develop novel chemotherapeutic agents.

## Discussion

DNA replication immortality is one of the critical hallmarks of cancerous cells that helps in continuous proliferation, invasion, and insensitivity to chemo- and radio-therapeutics (Swanton et al. 2024). The efficacy of cisplatin is compromised because of evolving intrinsic and acquired resistance mechanisms (Dasari and Tchounwou 2014). Resistance to cisplatin in various tumors has been reported to require TLS activities of Pol17, Rev1, and Polσ, with the former acting as an inserter and the others as mismatch extenders to maintain continuous replication and cell proliferation (Sharma et al. 2012; Zhao et al. 2012; Yoon et al. 2021). The roles of the other two TLS Pols-κ and τ, in cisplatin resistance is poorly established. This study revealed that Polκ is overexpressed exclusively in head and neck cancerous cells upon cisplatin treatment and also in cisplatin resistant oral squamous cell carcinoma cells. Other reports also suggested upregulation of Polκ in lung, ovarian, glioma, and prostate cancers, and its downregulation in stomach, colorectal, and breast cancers in comparison to the normal neighboring tissues associated with tumorigenesis, disease progression, and resistance to various therapeutics (Pillaire et al. 2014; Peng et al. 2016). According to the GCO (Global Cancer Observatory), the incidence of OSCC was 377,713 cases in 2020, which is believed to increase to 40% by 2040 (Tan et al. 2023). OSCC treatments require surgery followed by, if necessary, radio- or chemo-therapy. The chemotherapeutics include cisplatin, alone or in combination with 5-fluorouracil and docetaxel (TPF). But due to the development of chemoresistance, patients experience relapse, which leads to continued tumor growth and metastasis. Thus, understanding the mechanism of cisplatin resistance is key to identifying of additional targets and advancing drug discovery to improve OSCC management. We leveraged available cisplatin-resistant OSCC models to investigate the underlying mechanisms of Polκ’s involvement in cisplatin resistance and further suggested a few therapeutic targets (**Figure 11**).

**Figure 11:**
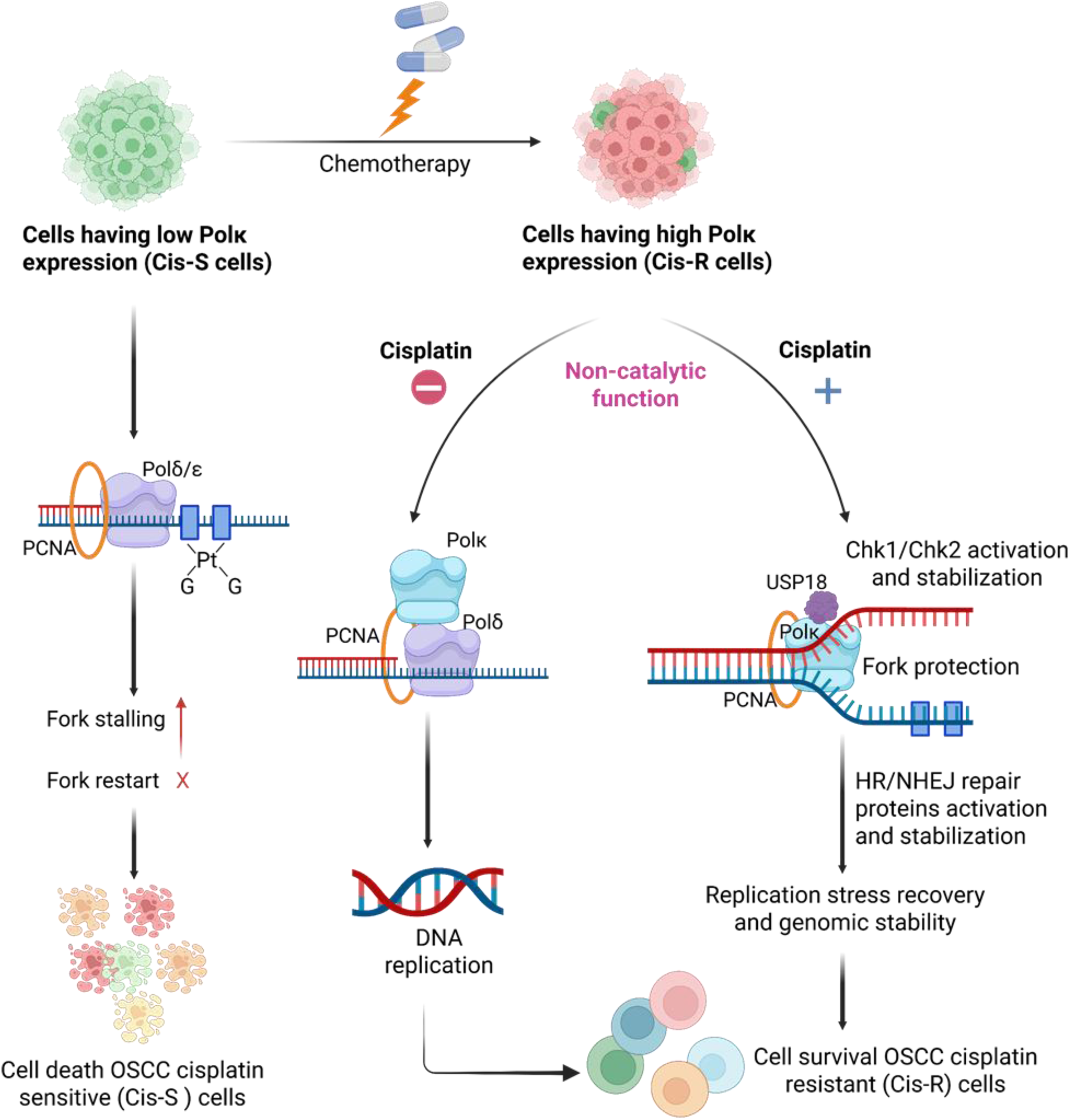
Dual tripartite interactions mediated by Polκ govern uninterrupted DNA replication and genome stability in OSCC cisplatin resistant cells. In cells having low Polκ expression, the bulky adducts generated by cisplatin could not be bypassed, leading to more fork stalling and collapse, which ultimately leads to cell death. Whereas in cisplatin-resistant cells where Polκ is overexpressed, it modulates two axes independent of its catalytic activity. The PCNA-Polκ-Polδ axis facilitates efficient replication and proliferation of cells; the other axis, PCNA-Polκ-USP18, stabilises DNA damage response proteins to activate checkpoint and HR/NHEJ repair pathways during cisplatin exposure. Both PIP and UBZ, but not the catalytic motifs of Polκ involved in PCNA and USP18 interactions, are indispensable for its role in cisplatin resistance.

### TLS independent function of Polκ stabilizes the cell’s genome to induce cisplatin resistance

The reversal to re-sensitization of resistant HNSCC cells to cisplatin due to the *POLK’*s absence suggested that Polκ could play an important role in the bypass of cisplatin adducts in these cells. Moreover, contrasting phenotypes of OSCC cells resulting from *POLK* depletion and overexpression in OSCC sensitive cells further underscored the critical role of Polκ in cisplatin adduct bypass and the minimal role of other TLS polymerases. Consequently, deletion of *POLK* led to OSCC-R cells with more DNA breaks and increased ψH2A.X expression, followed by a slowing of cell cycle progression, resulting in PARP-1 cleavage and enhanced cell death upon cisplatin exposure. The involvement of this polymerase in replication fork protection and recovery was again demonstrated in the DNA fibre assay. Further accumulation of stalled forks, fork conversions; and of slower moving ongoing forks, in *POLK* knockout cells than in cisplatin-resistant cells harboring abundant Polκ further strengthened the critical role of this TLS polymerase in reducing cisplatin toxicity. Interestingly, for such a function of Polκ, the catalytic activity was dispensable, but its PCNA interaction motifs were required. Thus, we concluded that the non-TLS of activity Polκ induces cisplatin resistance in OSCC. Yet another study reported that Polκ is overexpressed in melanoma, lung, and breast cancer cells, thereby promoting cellular proliferation in response to oncogenic alterations, specific kinase inhibition, oxidative stress, or starvation. Additionally, it enhances A375 melanoma cell’s resistance to the BRAF-targeted inhibitor vemurafenib; thus, targeting Polκ may reduce drug resistance in such cancer patients. The study even proposed a noncatalytic function of Polκ to explain the drug resistance (Temprine et al. 2020). Another report has also demonstrated a noncatalytic function of Polκ in DNA damage response (Kanemaru et al. 2015). Ectopically overexpressing catalytically dead Polκ mutant in human lymphoblastic Nalm6 cells rendered similar sensitivity to hydrogen peroxide and menadione as the wild-type cells, whereas the respective *POLK* knockout cells exhibited severe sensitivity, although the underlying mechanisms by which the catalytically deficient Polκ protected against such damage remained unexplored. However, another study suggested that Polκ induces ATR-CHK1 activation (Peng et al. 2016). Thus, Polκ’s non-catalytic activities are critical to multiple forms of cancer drug resistance across several tumor types.

### Polκ maintains cellular homeostasis of critical checkpoint and DNA break repair proteins

Next, we presented several compelling pieces of evidence supporting non-catalytic roles of Polκ in cellular homeostasis by regulating the abundance of critical proteins involved in checkpoint activation and DNA break repair pathways in human cells. For the first time, we showed that the depletion of Polκ in OSCC-R cells (both H357-R and SCC9-R) induces a decrease in the protein levels of Chk1, Chk2, p-Chk1, and p-Chk2 of ATM-ATR; CtIP, BRCA1, BRCA2, Rad51, and RAD54 of HR; and DNA-PKCs, Artemis, and DNA ligase IV of NHEJ pathways. Although we have not examined other proteins in these pathways, it appears that only the downstream proteins are affected by Polκ depletion. In the presence of a proteasome inhibitor, the protein levels of these recover in *POLK*-KO cells, again suggesting the role of Polκ in regulating the proteasomal degradation of these critical proteins. Conversely, in OSCC-R cells, Polκ overexpression maintains critical cellular abundance of these proteins, thereby protecting the replication fork and enabling further repair by these pathways to overcome cisplatin toxicity. DNA replication stress is nothing but slowing or stalling of replication fork progression, resulting in inhibition of DNA synthesis. The cellular response to replication stress requires activation of the ATR/Chk1 and ATM/Chk2 checkpoint axes, which sense the slow moving and stalled replication forks (Kumari et al. 2025). Their activation protects of the fork from nucleases such as MUS81 and MRE11 and enables selective repair pathways, prevents firing of late replication origins, and inhibits further progress to mitosis until replication is complete (Gonzalez Besteiro and Gottifredi 2015). Under normal physiological conditions, the ablation of Chk1 inhibits the replication rate. Accordingly, *CHEK1* knockout mice do not survive, and its haploinsufficiency enhances genome instability and mutagenesis (Stancel et al. 2009). During the course of this study, yet another report suggested that Chk1was stabilized specifically by Polκ, independent of its polymerase activity, in four different cell lines, including MRC5-SV, 293T, HCT116, and RKO (Dall’Osto et al. 2021). In that report, the expression levels of Chk2 and HR, and NHEJ pathway proteins were not examined. The replication fork restart defects associated with Polκ depletion can be rescued by Chk1 overexpression (Forment et al. 2011; Thompson et al. 2012). However, how Polκ protects Chk1 from proteasomal degradation was not determined until this study. The *CHEK1* expression at the transcriptional level is known to be p53 dependent (Gottifredi et al. 2001). Since the mRNA level of *CHEK1* remained unaltered in the Polκ knockout cells, whereas its protein expression was affected, it suggested that the regulation of Chk1 expression is p53 independent but Polκ dependent. Chk2 is a central transducer of the DNA damage response (DDR) that is activated by ATM upon DNA damage to halt cell cycle progression (at G1/S or G2/M phases) and to induce DNA repair or apoptosis (via p53). Chk2 also participates in various cellular functions in the absence of nuclear DNA lesions (Zannini et al. 2014). Upon activation, Chk2 phosphorylates nearly 24 nuclear proteins, including BRCA1, BRCA2, etc., that are involved in DDR (Seo et al. 2003). BRCA1 phosphorylation by Chk2 recruits Rad51 to lesion sites, promoting DNA strand invasion and the exchange steps of HR (Ciccia and Elledge 2010). Whereas BRCA2 phosphorylation causes disruption of the Rad51-BRCA2 complex, also allowing Rad51 to bind lesion sites (Bahassi et al. 2008; Saxena et al. 2019). No syndromes or cancer predisposition in mice have been linked to the absence of Chk2, although *CHEK*2^-/-^ mice are more susceptible to skin tumors (Hirao et al. 2002). NHEJ is the primary, relatively low-fidelity repair mechanism for double-strand breaks in mammals, active throughout the cell cycle but especially crucial in G1, and involves proteins such as Ku70/Ku80, DNA-PKCs, DNA Ligase IV/XRCC4, etc. While NHEJ directly repairs DNA double-strand breaks without a template, Chk2 mediates cell cycle arrest to allow sufficient time for this repair. Since some of these proteins in the ATR-Chk1, ATM-Chk2, HR, and NHEJ axes are stabilized in OSCC-R cells in a Polκ-dependent manner, we propose that while ATR-Chk1 is critical for replication stress management by preventing fork degradation and break accumulation without cisplatin, the ATM-Chk2 axis along with HR and NHEJ pathways may efficiently repair cisplatin adducts.

### A Polκ-Polδ-PCNA tripartite axis promotes efficient DNA replication of OSCC-R cells

Apart from TLS, the involvement of DNA polymerase kappa has also been implicated in the progression of replication forks through intrinsic barriers where replicative Pols have difficulty synthesizing DNA (Barnes et al. 2017). Common fragile sites (CFSs) are typically enriched in repetitive DNA sequences and are difficult to replicate; therefore, they are unstable genomic loci that are frequently altered in cancers. Biochemical assays revealed that Polκ replicate CFS-mimicking sequences more efficiently than Polδ. A recent study using Polκ sequencing revealed that Polκ activity is required for the replication of euchromatin regions, GC-rich regions, promoters, and DNA replicated early in the S phase of DNA replication (Tang et al. 2024; Torres et al. 2024). Polκ is also required to protect and restart replication forks following starvation for deoxynucleoside triphosphates (dNTPs), a process regulated by both the Fanconi Anaemia (FA) pathway and PCNA polyubiquitination (Tonzi et al. 2018). However, in this study, the involvement of catalytic activity of Polκ was not explored. Also, surprisingly, depletion of Polκ by siRNA did not alter Chk1 expression level. Probably, the use of different Polκ depletion strategies in different cell lines used in these studies might have resulted in these discrepancies. Nevertheless, all these reports suggested that Polκ’s role is not limited to damage sites and that it plays multiple cellular functions as well. The colony counting, Comet and fibre assays, cell cycle analysis, and cell survival of OSCC-R cells in the presence of replication inhibitors like HU, Aphidicolin, and Camptothecin in this study further strengthen the role of Polκ in regulating replication stress tolerance and fork progression. However, contrary to the *in vitro* DNA replication assay using the CFS template, we observed that in this context as well, the catalytic activity of Polκ is not required. A cooperative tripartite interaction among Polκ, PCNA, and Polδ appears critical for the high proliferative capacity of OSCC-R cells. In fact, studies have shown that the depletion of Polκ is associated with increased rates of spontaneous mutagenesis and inflammation in mice (Stancel et al. 2009; Hakura et al. 2019). Thus, Polκ plays a critical role in minimizing replication stress by stabilizing Polδ, or possibly other replicative DNA Pols, on the replication fork and by activating the ATR-Chk1 pathway to protect stalled replication forks at difficult replication sites.

### PCNA-Polκ-USP18 axis promotes efficient replication of cisplatin adducts in the genome of OSCC-R cells

So far, we have established the role of Polκ in genomic DNA replication without genotoxic stress and in protecting replication forks by activating ATR-Chk1, ATM-Chk2, HR, and NHEJ pathways, thereby, regulating their protein stability and causing cisplatin resistance. Ubiquitylation and deubiquitylation play important roles in regulating DNA repair pathways by coordinating the recruitment and removal of repair proteins at and from the damaged site (Messick and Greenberg 2009; Borsos et al. 2020; Sharma et al. 2020; Tang et al. 2021). Reports suggested that Chk1 function is regulated by ubiquitination by the cullin-ring E3-ubiquitin ligases CUL1, CUL4A, and HUWE1 conjugating systems and deubiquitination by ubiquitin hydrolases like USP1, USP3, USP7, and ataxin3, which deubiquitinate Chk1 (Zhang et al. 2005; Zhang et al. 2009; Guervilly et al. 2011; Alonso-de Vega et al. 2014; Tu et al. 2017; Cheng and Shieh 2018; Cassidy et al. 2020). Similarly, Chk2 ubiquitination is controlled by Cullin 1, PIRH2, and SIAH2, and deubiquitination by USP7 and USP39 (Lovly et al. 2008; Bohgaki et al. 2013; Garcia-Limones et al. 2016; Wu et al. 2019; Liu et al. 2024). E3 ligases like RNF8, RNF168, and BRCA1 are crucial for post-translational modification that regulate DNA double-strand break (DSB) repair by recruiting repair factors, such as 53BP1 and BRCA1, to damage sites. Specific E3 ligases, such as RFWD3 and RNF4, mediate the ubiquitin-mediated degradation of RPA and Rad51 proteins to regulate their turnover during homologous recombination. At least 15 of USPs (USP1, USP10, USP11, USP15, USP20, USP26, USP29, USP3, USP37, USP4, USP42, USP44, USP6, USP7, and USP51) have been demonstrated to regulate DSBR. These USPs can promote or suppress the choice of a DSBR pathway. Our transcriptomic analysis of OSCC-R RNA-seq data of revealed upregulation of certain DUBs and downregulation of proteins involved in ubiquitin-mediated proteasomal degradation. We proposed that preventing the degradation of key DDR proteins could enable OSCC-R cells to efficiently nullify the cisplatin effect. Although we could not directly assess Polκ’s role in the downregulation of 26S proteasomal proteins, we showed a direct association of Polκ with the deubiquitinase USP18 via its UBZ domain, thereby regulating a critical mass of proteins affected in the absence of Polκ. How Polκ absence alters even the USP18 transcripts required further investigation. For the first time, we reported a PCNA-Polκ-USP18 deubiquitinating system in replication fork protection and restart, which is responsible for chemoresistance. PCNA bound Polκ recruits USP18 to the replication fork via its UBZ domain forming a tripartite axis that stabilizes DDR proteins. This study has also ruled out a direct interaction between PCNA and USP18, as supported by the lack of a bonafide PIP motif in the USP18’s primary sequence. The PIP and UBZ domain mutants of Polκ render phenotypes similar to those of complete deletion of Polκ or USP18, suggesting the importance of this axis in cisplatin resistance. A recent study already demonstrated the role of USP2, which deubiquitinates NBS1 and increases the stability of the MRN complex (MRE11–RAD50–NBS1) during the DSB response (Kim et al. 2023). However, USP2 may function independently of Polκ, as we did not see any physical interaction between the two proteins. Several studies have highlighted the importance of ubiquitination/deubiquitination systems in governing tumor behaviors including proliferation, migration, invasion, and resistance to therapy (Sharma and Almasan 2020; Chen et al. 2022; Zhang et al. 2023). Along similar lines, this study found that USP18 depletion reversed of the chemoresistant phenotype in OSCC-R cells.

### Implications of Polκ as a chemotherapeutic target

Due to the central roles of TLS Pols in mutagenesis and cell survival post DNA damage, they have emerged as potential targets for the development of anticancer drugs that can be used alone or in combination with existing therapies. For example, small molecule inhibitors against human Rev1 have been designed to target its interacting surface with Polσ and other TLS Pols, and some are being considered for clinical trials (Dash et al. 2018; Wojtaszek et al. 2019; Chatterjee et al. 2020; Yoon et al. 2021). However, targeting Rev1 for cancer treatment may not be viable, as it will inhibit TLS across all types of lesions in normal cells and increase cytotoxicity and tumorigenicity. Although Pol17 and Polσ are involved in cisplatin resistance, suitable inhibitors against these polymerases are yet to be identified and validated as chemotherapeutics (Korzhnev and Hadden 2016; Patel et al. 2021). This study unveils multiple non-conventional roles of Polκ in replication fork stabilization, efficient replication restart, checkpoint activation, and cellular homeostasis of critical DSBR proteins, thereby, inducing cisplatin resistance by multiple axes. Since the catalytic activity of Polκ is dispensable for most of these phenotypes, a specific small molecule that can block its interaction with PCNA, rather than an existing generic inhibitor T2AA, can be designed and explored as adjunct therapy. Additionally, effective USP18 inhibitors can also be explored as additional therapeutics.

## Materials and Methods

### Cell lines and cell culture

Various human transformed cell lines such as A549 (non-small cell lung cancer), HEK293 (embryonic kidney), MCF7 and MDA-MB-231 (breast cancer), MIA PaCa-2 (pancreatic cancer), SH-SY5Y (Neuroblastoma), DU145 (prostate cancer), H357 and SCC9 (tongue OSCC) and Huh-7 (hepatoma) are maintained and grown appropriately, and used regularly in our laboratory (Khandagale et al. 2019; Sahu et al. 2024; Sahu et al. 2026). The cisplatin resistant OSCC counterparts were developed in Dr Rupesh Dash’s laboratory (Maji et al. 2019). Briefly, A549, HEK293, MCF7 and MDA-MB-231, SH-SY5Y, Huh-7, MIA PaCa-2 and DU145 cells were cultured in Dulbecco’s Modified Eagle Medium (DMEM) from Hi-Media (Cat#AL007A). H357, SCC9 cell lines were cultured in Dulbecco’s Modified Eagle Medium / Nutrient Mixture F-12 Ham [DMEM / F12, 1:1] from Hi-media (Cat#AL139A). All culture media were supplemented with 10% FBS (Gibco, Cat# A5256701), 1% penicillin-streptomycin (PS from Gibco, Cat#15140122), except for SH-SY5Y, where 15% FBS was added. For Huh7 and SH-SY5Y culturing, additionally, MEM non-essential amino acids (Gibco, Cat# 11140-050) were supplemented. Similarly, 0.5μg/ml sodium hydrocortisone (Sigma, Cat# H6909) was added to the medium of the H357/SCC9 culture. All cell lines were kept in a 37°C incubator with 5% CO2 and 95% relative humidity.

### Drug supplements, oligonucleotides and antibodies

cis-Diammineplatinum(II) dichloride (Cat#P4394), T2AA (Cat#SML0794), and Camptothecin (Cat#C9911) were purchased from Sigma-Aldrich. MG132 (Cat#A2585) was sourced from ApexBio. Aphidicolin (Cat#38966-21-1) and hydroxyurea (Cat#H0310) were procured from MP Biomedicals and TCI, respectively. The list of primers used in this study is provided in Table 1, and the antibodies are listed in Table 2.

**Table 1:**
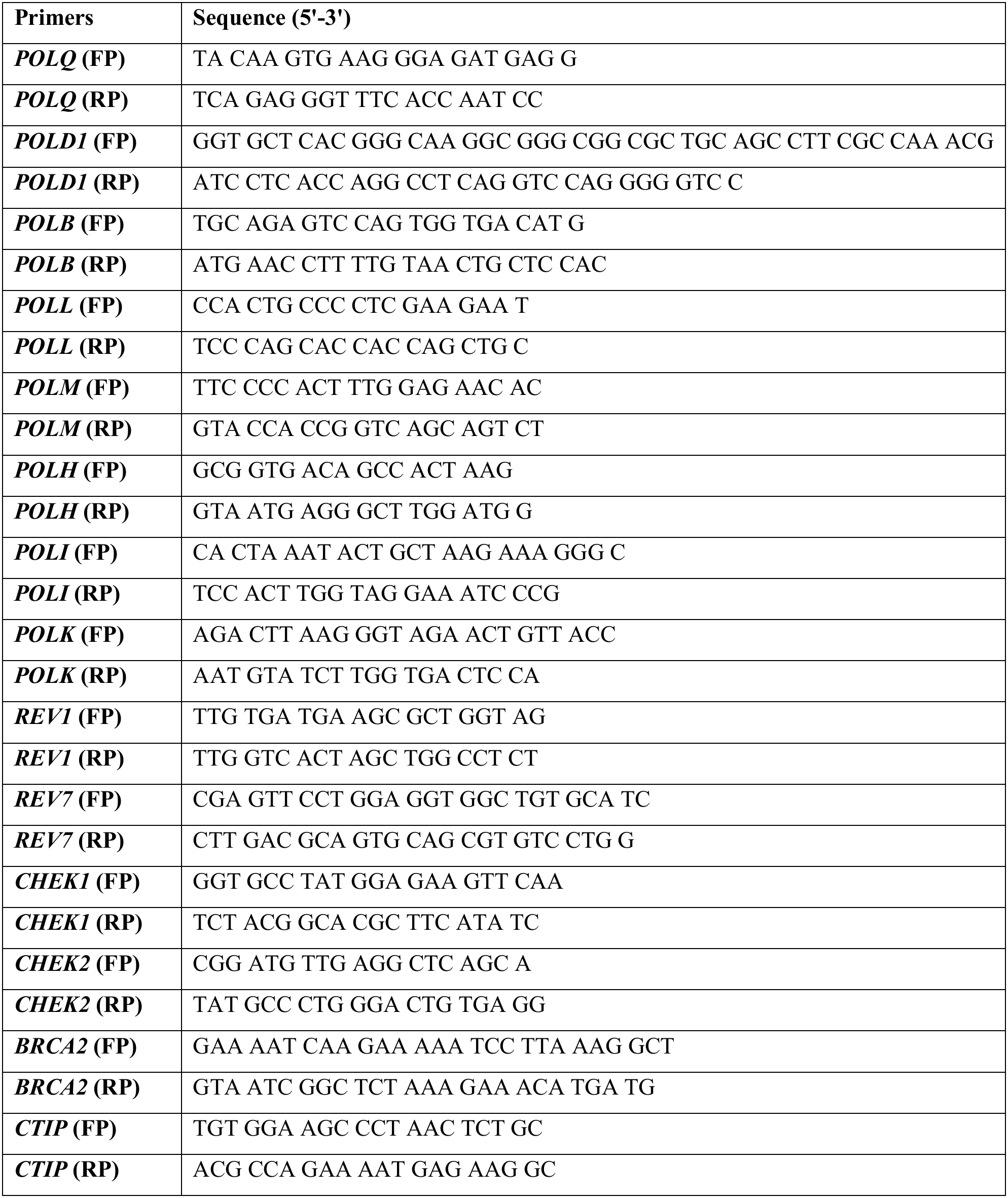
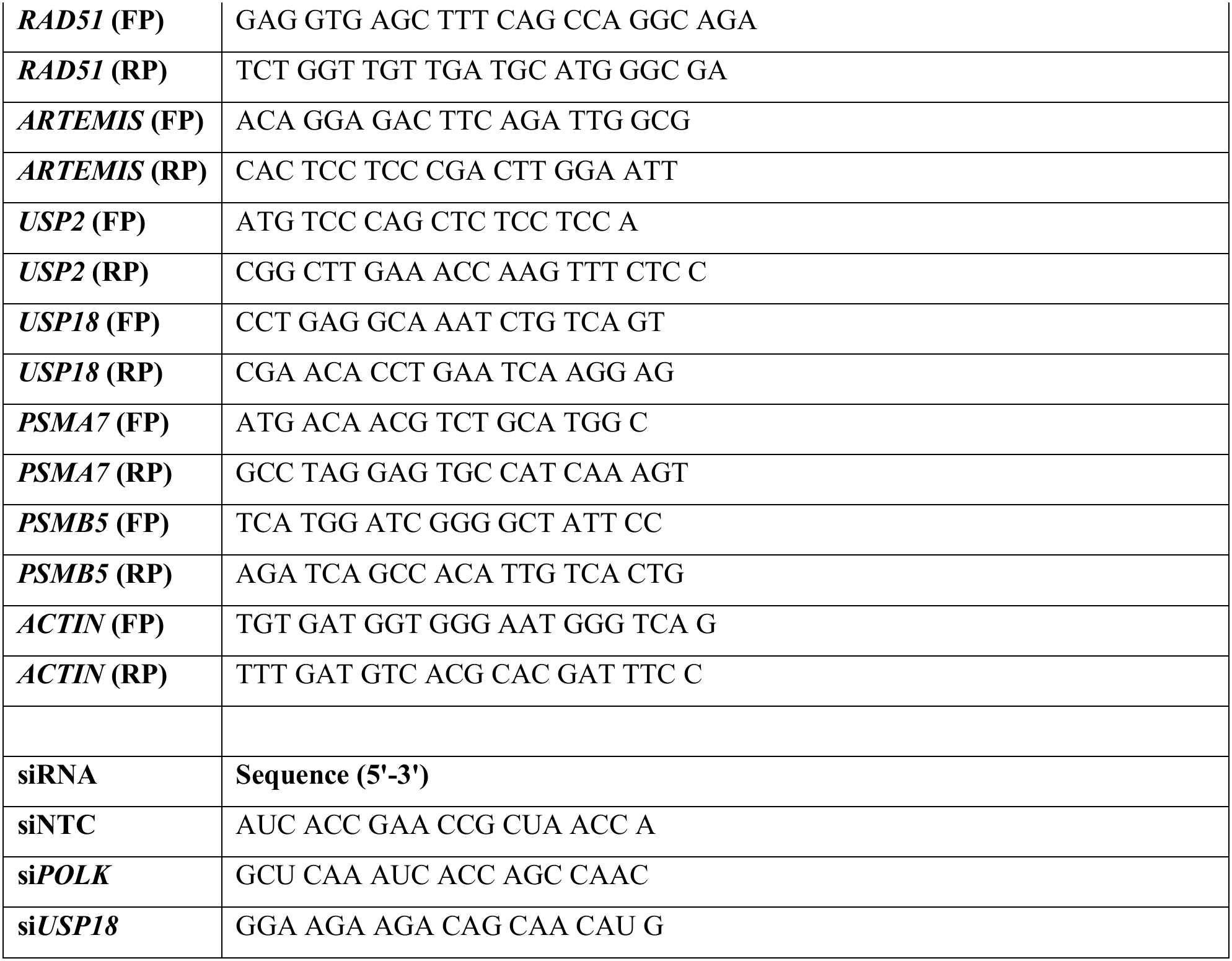
List of oligonucleotides used in the study.

**Table 2:** List of antibodies used in the study.

| **Antibodies** | **Source** | **Identifier** |
|---|---|---|
| Polκ | Santa Cruz | sc-166667 |
| Polη | CST | 13848S |
| Polθ | Self-raised | n.a. |
| P125 | Sigma | SAB4200053 |
| Polβ | Santa Cruz | sc-376581 |
| Polλ | Santa Cruz | sc-373844 |
| Polμ | Santa Cruz | sc-398666 |
| Polι | Santa Cruz | sc-101026 |
| Rev1 | Santa Cruz | sc-393022 |
| Rev7 | Santa Cruz | sc-47747 |
| β-actin | Santa Cruz | sc-47778 |
| Chk1 | Santa Cruz | sc-8408 |
| Rad51 | Santa Cruz | sc-53428 |
| γH2AX (S139) | CST | 9718S |
| PARP1 | CST | 9542S |
| GFP | CST | 2956S |
| ATR | CST | 2790S |
| ATM | CST | 2873S |
| P-ATR (Ser428) | CST | 2853S |
| P-ATM (Ser1981) | CST | 5883S |
| Chk-1 | CST | 2360S |
| Chk-2 | CST | 2662S |
| p-Chk1 (S341) | CST | 2348S |
| p-Chk2 (T68) | CST | 2661S |
| Mre11 | CST | 4895S |
| Rad50 | CST | 3427S |
| NBS1 | CST | 3002S |
| P-NBS1 | CST | 3001S |
| CtIP | CST | 9201S |
| BRCA1 | CST | 14823T |
| BRCA2 | CST | 10741T |
| RAD54 | CST | 15016T |
| Rad51 | CST | 8875T |
| KU70 | CST | 4588T |
| KU80 | CST | 2180T |
| DNA PKCs | CST | 38168T |
| P-DNA PKCs | CST | 68716T |
| Artemis | CST | 13381T |
| DNA LIG IV | CST | 14649T |
| XLF | CST | 2854T |
| XRCC4 | CST | 23908S |
| Histone H3 | CST | 14269S |
| Cleaved PARP1 (Asp214) | CST | 9541S |
| p50 | Sigma | SAB4200054 |
| p12 | Self-raised | n.a. |
| Ki 67 | CST | 12202 |
| USP2 | Proteintech | 10392-1-AP |
| USP18 | CST | 4813S |
| PSMA7 | Proteintech | 15219-1-AP |
| PSMB5 | CST | 12919S |

n.a.: Not applicable

### Generation of *POLK* knockout cell lines

*POLK* gene knockout in H357-R cells was achieved using a CRISPR/Cas9 gene-targeting approach as recommended by Santa Cruz Biotechnology. Cells were co-transfected with the human *POLK* CRISPR/Cas9 KO plasmid (Cat#sc-405052) and its HDR plasmid (Cat# sc-405052-HDR) using Viafect transfection reagent (Promega, Cat#3498). The CRISPR plasmid pool contained three individual guide RNAs along with a Cas9 nuclease expression gene, while the HDR plasmid facilitated homologous recombination repair at the targeted loci, introducing a Red Fluorescent Protein (RFP) gene and a puromycin resistance gene. Transfectants were subjected to selection in a medium containing 2μg/ml puromycin for 48 hours. A single cell was selected by fluorescence-activated cell sorting (FACS), placed in 96-well plates, and cultured for 7-10 days to allow the development of individual colonies. Selected knockout clones were confirmed by Western blotting to verify successful *POLK* gene knockout.

### siRNA transfection and gene knockdown

Cells were seeded for 24 hours before transfection with 25nM siRNA (Eurofins) using Lipofectamine 2000 (Invitrogen). The following siRNA molecules were used: siNTC: 5′-AUC ACC GAA CCG CUA ACC A-3′, si*POLK*: 5′-GCU CAA AUC ACC AGC CAAC-3′, si*USP18*: 5′-GGA AGA AGA CAG CAA CAU G-3′. After 48 hours of transfection, cells were harvested for further experiments.

### Generation of plasmid constructs and over-expression

Amino terminal GFP tag constructs of wild type Polκ and its PIP (F532A, L533A, F868A, F869A), catalytically deficient (D198A, E199A) and UBZ ( C624A, C627A, C779A, C782A) mutants were generated and designated as GFP-WT, GFP-PIP, GFP-CD, and GFP-UBZ, respectively. To generate the PIP mutant, the C-terminal portion harboring both mutated PIP motifs was amplified using 5′-CC CGG ATA TCT AGT TTT CCC AAT GAA GAG GAC AGG AAA CAC CAA CAA AGG AGC ATT ATT GGC GCT GCA CAG GCT GG-3′ and 5′-GGCC AAG CTT GGA TCC TTA CTT GCA GCA TAT ATC AAG GG-3′ primer pairs, digested with EcoRV and HindIII restriction enzymes and ligated into the same sites to replace the wild type fragment in pCMV-human Polκ. Similarly, a ∼2 kb fragment with a catalytically dead mutation was amplified using primer pairs 5′-TTT TAT GGC CAT GAG TCT TGC CGC GGC CTA CTT G-3′ and 5′-AGC GGA TAA CAA TTT CA-3′, digested with MscI and HindIII, and cloned into the same sites to replace the wild-type fragment in pUC19-human Polκ. To generate a UBZ mutant, the C-terminal portion of Polk was amplified using primer pairs 5′-CA GAG AAT TCA GAT GAC TGT CAG ATA CTT ACC GCT CCT GTT GCC TTT AGG-3′ and 5′-GGT TAG ATC TGA AGT CTT TTG TTC TAC GTT AGC AAC AGG AGC AAC TAG AGC-3′, digested with EcoRI and BglII, and ligated to replace similar wild type fragment from a modified pUC19-Polκ construct that lacks the EcoRI site of multiple cloning site. BamHI fragments of these constructs were subcloned into the same site of pcDNA-GFP. Various constructs of WT and mutants of Polκ were transfected into H357-R *POLK* KO cells, H357-S, and SCC9-S cells using FuGENE HD transfection reagent (Promega, Cat#E2311) according to the manufacturer’s instructions. After 48 hours of transfection, the cells were harvested for further experiments.

### RNA extraction and RT-PCR

Total RNA was purified using TRIzol (Ambion Life Technology, Cat#15596018). After purification, quality and concentration of RNA were checked using a NanoDrop 2000, and cDNA was generated using an Applied Biosystems cDNA synthesis kit (Cat#4368814). The required genes of interest were amplified by RT-PCR; conditions included denaturation at 95 °C, annealing at 55 °C, and extension at 72 °C. Bands were visualised by agarose gel electrophoresis; photographs were captured with a Bio-Rad ChemiDoc MP imaging system and analysed with ImageJ software. For real-time quantitative PCR, all reactions were performed in technical duplicates and biological triplicates by amplifying the gene of interest and *ACTIN* as an endogenous control using SYBR Green Master Mix (Applied Biosystems, Cat#A25742) and QuantStudio 3 (Applied Biosystems). The ΔΔct values were calculated and compared. Graphs were plotted using GraphPad Prism 9.0.0 software after calculating the fold change.

### Western blot analysis

Cells with 70-80% confluency were collected via trypsinisation and then lysed on ice for 30 minutes with RIPA lysis buffer (CST, Cat#9806), a protease inhibitor cocktail (BioPioneer, Cat#BPBOI001), and a phosphatase inhibitor cocktail (CST, Cat#5870S) with quick vortexing every 5 minutes. Samples were centrifuged at 14,000 g for 20 minutes at 4℃, and supernatants were collected. Total protein concentration was measured using the Bradford assay. For nuclear extracts, a protocol from Thermo Fisher Scientific, with slight modifications, was used. Briefly, cells (2×10^6^) were collected and washed twice with ice-cold 1× PBS. The cell pellet was resuspended in 200 μl hypotonic solution (20 mM Tris-HCl (pH 7.4), 10 mM NaCl, 3 mM MgCl2) by pipetting up and down multiple times. Samples were kept on ice for 15 minutes, followed by the addition of 10 μl of 10% NP-40 and vortexing for 10 seconds at high speed, then centrifuged for 10 minutes at 5,000 rpm at 4℃. Supernatants were collected and saved as the cytoplasmic fraction. The purity of the cytoplasmic fraction was confirmed by checking the expression of GAPDH, as it is present in abundance in the cytoplasmic fraction and absent or minimal in the nuclear fraction. The pellet was then resuspended in 40 µl of complete cell extraction buffer with high salt concentrations (400 mM NaCl). After 30 minutes on ice with vortexing every 5 minutes, samples were centrifuged at 14,000 g for 20 minutes at 4℃. Supernatants were collected and saved as nuclear fractions, and after protein estimation using Bradford’s assay, about 40 μg of total protein from each sample was boiled in SDS buffer at 95°C for 10 minutes, resolved on 7-12% SDS-PAGE gels, and transferred to polyvinylidene difluoride (PVDF) membranes (Cytiva, Cat#10600023). To prevent non-specific binding, membranes were blocked with 5% non-fat dry milk (NFDM, Hi-Media, Cat #GRM1254) or BSA (MP Biomedicals, Cat#160069) in TBS-T buffer [Tris-Buffered Saline with 0.1% Tween20 (Gene Tex, Cat#GTX03634)] at room temperature for 1.5 hours. Blots were then incubated with specific primary antibodies against various proteins at 4°C overnight on a rocking platform. After three washes with TBS-T, membranes were incubated for 1.5 hours at room temperature with HRP-conjugated secondary antibodies (1:5000) (anti-rabbit, Cat#A0545, anti-mouse Cat#A9044 and anti-rat Cat#A9037) from Sigma. Protein bands were visualised using a chemiluminescent HRP substrate ECL reagent (TaKaRa, Cat#T7101A) on a Chemi-Doc MP imaging system (BIO-RAD). Protein expression was quantified using ImageJ software.

### Immunoprecipitation

Various cell types were grown in 100-mm tissue culture dishes. After 24 hours, cells were collected by trypsinisation and lysed in 400 µl of 1X immunoprecipitation (IP) lysis buffer from the Protein G immunoprecipitation kit (Sigma, Cat #IP50), containing Protease/Phosphatase Inhibitor Cocktail, on ice. For overexpressed sets, cells were transfected with GFP-WT, GFP-PIP, and GFP-UBZ plasmids using a previously described transfection method, and cells were collected after 48 hours; similar steps were followed. The lysate was centrifuged at 14,000 g for 20 minutes at 4°C. Supernatant protein concentration was determined using Bradford reagent (Sigma, Cat#B6916). Freshly prepared samples containing 500 µg-1 mg of total protein were pre-cleared via incubation with protein G agarose beads (Sigma, Cat#IP50) for 4 hours at 4°C on a rotator and used for IP by incubating with anti-Polκ antibody, anti-PCNA, anti-USP2, anti-USP18, anti-Polθ or IgG control and 1X IP buffer to make up the final volume to 600 µl overnight at 4°C on a rotator. Immune complexes were then precipitated by protein G agarose beads at 4°C overnight. The next day, the immunoprecipitates were washed 6 times with 700 µl 1X IP lysis buffer on ice. The last wash was done with 0.1X IP buffer. An empty spin was performed for 30 secs to remove any remaining buffer. Then samples were resuspended in 40 µl of 1X Laemmli sample buffer. The samples were separated by 10% SDS-PAGE, and Western blot analysis was performed as described. Immunodetection was done using primary p50, p12, PCNA, Polκ, USP2, USP18, and GFP antibodies, and respective HRP-conjugated secondary antibodies (anti-mouse Cat#A9044, anti-rabbit Cat#A0545, anti-rat Cat#A9037) from Sigma. ECL reagent (TaKaRa, Cat#T7101A) was used for image development on the Chemi-Doc MP system (BIO-RAD). Whole-cell lysate (input) was also subjected to immunoblotting to detect the same protein expression as described.

### Colony-forming assay

In the clonogenic survival studies, approximately 500 cells were seeded into each well of a 6-well plate or a 35-mm Petri plate. After 24 hours, cisplatin was administered at the indicated concentrations for 48 hours. After 48 hours, the media was replaced with complete growth media. Then cells were allowed to grow for a further 11–14 days; media changes were done at 2-day intervals. The surviving colonies were fixed with 4% formaldehyde, treated with crystal violet (0.5%), and images were captured with the BIO-RAD Chemi-Doc system and analysed in ImageJ.

### Annexin-V/PI assay

Cells were cultured in 6-well plates for 24 hours and subsequently treated with cisplatin (5 µM). The Annexin V/PI flow cytometry assay kit (BD Biosciences, Cat#556547) was employed to assess cell death in H357-S/R and H357-R *POLK* KO cells. Cells were harvested 48 hours post-treatment via trypsinisation. Following the manufacturer’s guidelines, the cells were stained with Annexin V and Propidium Iodide (PI) after a single wash with 1X PBS. In brief, 100 μl of 1X annexin V binding buffer was added to the cells, which were subsequently incubated in the dark for 15 minutes with FITC-conjugated annexin V to assess phosphatidylserine translocation, alongside Propidium iodide (PI) to evaluate membrane integrity. Ultimately, each reaction was adjusted with 400 μl of 1X annexin V binding buffer just before the flow cytometry analysis (BD LSR Fortessa). Apoptosis percentages were quantified using FlowJo V.10 software.

### Immunofluorescence staining

Cells were seeded at 70% confluency on glass coverslips under standard cell culture conditions and allowed to attach for 24 hours. First, cells were fixed using 4% formaldehyde for 15 minutes and subsequently permeabilised with 0.2% Triton X-100 in PBS for 10 minutes. Cells were subsequently blocked using 10% FBS in 1X PBS with 0.2% Triton X-100 for 45 minutes to 1 hour at 37°C. The fixed cells were incubated overnight with primary antibodies (1:200-400 dilution) like Polκ, phospho-Histone H2A.X (Ser139), Ki-67, p12, p50, Polθ, USP2, USP18, Chk1, Chk2, P-Chk1, P-Chk2, BRCA2, Rad51, and Artemis. Following three washes with 1X PBS, the coverslips were incubated with secondary antibodies anti-mouse IgG Alexa Fluor 488 (Invitrogen, Cat#A-11017), anti-rabbit IgG Alexa Fluor 594 (Invitrogen, Cat#A-21207) and anti-rat IgG Alexa Fluor 555 (Invitrogen, Cat#A-21434) at a 1:500 dilution for 1 hour. The coverslips were further washed, stained with DAPI, and mounted. Images were obtained using a confocal laser scanning microscope (Stellaris 5, Leica, Germany). The foci were quantified using ImageJ software. The co-localization percentage and scatter plot were obtained from Stellaris 5. The percentage of Ki-67 positive cells was calculated as: (number of positive cells in a field/total number of cells in a field) X 100.

### Alkaline comet assay

Cells were seeded in 6-well plates and treated with cisplatin (5 µM) for 48 hours. Both control (i.e., untreated) and cisplatin-treated cells were trypsinised and collected. The cells were then mixed with 0.5% low-melting agarose (LMA, cat#1613111) and fixed onto slides pre-coated with 1.5% agarose. Lysis was carried out in 20-30 ml of lysis buffer (2.5 M NaCl, 0.1 M EDTA, 10 mM Tris-HCl; pH=10) for 1 hour at 4°C in the dark, followed by neutralization with a buffer (300 mM NaOH, 1 mM EDTA; pH>13) for 20 mins at 4°C. Electrophoresis was performed in an alkaline buffer for 30 mins at 20V, after which the slides were dried and stained with SYTOX Green. Images were captured using an Apotome microscope (Zeiss). Data analysis was performed using Open-Comet software.

### MTT cell viability assay

The colourimetric cell viability assay using 3-(4,5-dimethylthiazol-2-yl)-2,5-diphenyltetrazolium bromide (MTT) was performed in 96-well microplates. Cells (5×10^3^) were seeded in each well of a 96-well plate and then treated with the indicated concentrations of cisplatin and T2AA for 48 hours, and with HU, aphidicolin, and camptothecin for 24 hours. After treatment, the medium was removed, and MTT (0.5 mg/ml) in PBS solution was added directly into the wells and incubated for 3 hours. The medium was subsequently removed, and solvent (DMSO) was added to the cells to solubilize formazan crystals. The absorbance was measured at 570 nm after 20 minutes of shaking using a Multimode plate Reader (PerkinElmer).

### Cell cycle synchronization and cell cycle analysis

To arrest H357-S, H357-R and H357-R POLK KO cells in G1/S phase, cells were seeded into 10 cm2 cell culture plates (corning) at a confluency of 30-40% in complete growth media and then allowed to grow overnight at 37°C with 5% CO2. The next day, cells were treated with 2 mM thymidine in complete medium for 12 hours (initial synchronisation). Then cells were washed three times with 3 mL of 1X PBS and grown in a thymidine-free medium (first synchronisation release) for 15 hours, then washed three times with PBS and re-cultured in a complete growth medium containing 2 mM thymidine (second synchronisation) for another 12 hours. Finally, cells were washed three times using 3 mL of 1X PBS and released from the G1/S arrest. Cells were taken at 0, 3, 6, 9, and 12 hours. For cisplatin-treated sets, cisplatin was added to the medium during the second synchronisation release, and cells were collected at the specified intervals for cell cycle analyses. In brief, the cells were collected via trypsinisation and then washed twice with 2 ml of ice-cold 1X PBS. Then fixed in chilled 70% ethanol. After washing with 1X PBS, the cells were resuspended in a buffer containing 0.2% Triton X-100, 50 μg/ml propidium iodide (PI), and 100 μg/ml RNase A, and incubated for 30 minutes in the dark. The data were recorded with a minimum of 10,000 events on a BD LSR Fortessa and analysed using FlowJo V.10 software.

### DNA fibre assay

A standardised protocol for the DNA fibre assay to follow replication fork movement was followed. Briefly, cells were seeded for 24 hours prior to the start of the experiment. Then, cells were pulse-labelled with 30 μM CldU (Sigma, Cat#C6891) and 300 μM IdU (Sigma, Cat#I7125). For cisplatin-treated sets, cells were exposed to 5 μM cisplatin for 24 hours before adding CldU and IdU. In the HU treatment set, cells were exposed to 2 mM hydroxyurea for 2 hours between two pulses. Cells were collected and resuspended in ice-cold PBS at a density of 2.5 × 10^5^ cells/ml. Labelled cells were then mixed with unlabelled cells at a 1:1 ratio. Subsequently, 2.5 μl of the cell suspension was placed onto a clean Superfrost Plus microscope glass slide (Fisher Scientific, Cat#1255015). Cells were treated with 7.5 μl of lysis solution (200 mM Tris-HCl; pH 7.4, 50 mM EDTA, 0.5% SDS) and lysed by gently swirling with a 10 μl micropipette tip, then pipetting the mixture up and down. The slides were kept in parallel position for 8-10 minutes for lysis, then tilted to 45° to facilitate fibre spreading and allowed to air-dry for approximately 30 minutes. The samples were fixed in methanol and acetic acid solution (3:1) overnight at 4°C, washed twice with 1X PBS the following day, and then denatured with hydrochloric acid (2.5 N) for 1 hour. Subsequently, slides were incubated in a blocking solution (2% BSA, 0.1% Tween-20 in 1X PBS, sterile-filtered) and stained with anti-rat CldU antibody (1:200, Abcam, Cat#ab6326) and anti-mouse IdU antibody (1:100, BD Biosciences, Cat#347580) for 2.5 hours. Then the slides were subjected to five washes with PBS-T (0.2% Tween-20) for three minutes each. Subsequently, they were briefly incubated with a blocking solution (2% BSA, 0.1% Tween-20, and 1X PBS). Slides were treated with anti-mouse Alexa Fluor 488 (Invitrogen, Cat#A11001) and anti-rat antibody (Jackson ImmunoResearch, Cat#712-166-153). Following secondary antibody staining, the slides were washed with PBS-T five times for three minutes each. The slides were thoroughly air-dried and mounted using Fluoromount aqueous mounting media (Sigma, Cat#F4680), with coverslips sealed with clear nail polish. Slides were captured at 63X magnification using Stellaris5 (Leica, Germany), and fibre length was assessed using ImageJ software, and fork speed was calculated using the following formula: Average length of (IdU+CldU) per 30 mins (i.e., Time of pulse).

### Quantification and statistical analyses

Data were represented as mean ± SEM. Statistical analyses were performed using GraphPad Prism 9.0.0, and p-values < 0.05 were considered statistically significant.

### Availability of data and materials

Data sharing is not applicable to this paper.

## Authors’ contributions

Ipsita Subhadarsini: Writing – original draft, Validation, Investigation, Formal analysis, Data curation, Resources, Methodology, and Writing—review and editing; Jugal Kishor Sahu: Writing – original draft, Validation, Investigation, Formal analysis, Data curation, Resources, Methodology, and Writing—review and editing; Shweta Thakur: Writing – original draft, Validation, Investigation, Formal analysis, Data curation, Resources, Methodology, and Writing—review and editing; Rupesh Dash: Resources, Supervision, and Writing—review and editing; Narottam Acharya: Writing – original draft, Formal analysis, Conceptualization, Supervision, Funding acquisition, Project administration, and Writing—review and editing

## Acknowledgements

We thank Mr. Sitendra Prasad Panda, Mr. Bhabani Shankar Sahoo, and Mr. Paritosh Nath for their technical assistance in the laboratory, Imaging facility and FACS facility, respectively. Our laboratory colleagues are acknowledged for their thoughtful discussion. JKS and IS are CSIR-Senior Research Fellows, and ST is thankful to DBT-RA fellowship.

## Abbreviations

TLS: Translesion Synthesis
Pol: DNA polymerase
Polδ: DNA polymerase delta
PCNA: Proliferating cell nuclear antigen
DDR: DNA damage response
PIP: PCNA interacting protein
NER: Nucleotide excision repair
DSBs: Double strand breaks
HR: Homologous recombination
NHEJ: Non-homologous end joining
OSCC: oral squamous cell carcinoma
HU: hydroxyurea

