## Supplementary material for "PCNA-Polκ-Polδ/USP18 axes stabilize replication fork and restart to reduce cisplatin cytotoxicity": Supp. Text

<u>**Supplementary Information**</u>

**Supplementary Figure 1:** **(A)** Bar graphs showed differential expression of various DNA polymerases in terms of mRNA (**i**) and proteins (**ii**) in OSCC cells (H357-S, H357-R, SCC9-S, SCC9-R). Data represented as mean ± SEM (n=2), two-way ANOVA with Tukey's multiple comparisons test (*p < 0.05, and ****p < 0.0001). **(B)** Western blot analysis confirmed *POLK* deletion in H357-R cell lines. **(C)** Bar graph showed the fold change in the expression of Polκ when normalized to actin in H357-S, H357-R and H357-R *POLK* KO cells without and with cisplatin treatment, mean ± SEM (n=2). Statistical significance as determined by Ordinary one-way ANOVA, Dunnett's multiple comparisons test, (***p < 0.001, and ****p < 0.0001). **(D)** Representative immunofluorescence images of H357-S, H357-R and H357-R *POLK* KO cells revealed the expression of Polκ (green) puncta in nucleus (DAPI, blue), and their corresponding merged images in presence and absence of cisplatin (**i**). Quantification of Polκ foci numbers is indicated in a bar diagram as mean ± SEM (n=3) (**ii**). Statistical significance as determined by Ordinary one-way ANOVA, Tukey's multiple comparisons test, (****p < 0.0001, and ns- nonsignificant). Scale bars indicate 7.5 μm. **(E)** H357-S and SCC9-S cell lines were transfected with NTC siRNA and *POLK* siRNA as described in Methods. Lysates were prepared and Western blotting was performed with indicated antibodies. β-actin was used as an internal control. Representative immunoblots from three independent experiments are shown. Densitometric fold change relative to proteins in H357-S and SCC9-S cells has been provided just below each band (**i**) and (**ii**). **(F)** Cells were treated with the indicated concentrations of cisplatin drug for 48 hours and then an MTT assay was performed. The graph indicates the mean ± SEM (n=3), Statistical significance determined by two-way ANOVA, Šídák's multiple comparisons test, (**p < 0.01, ***p < 0.001, and ns- nonsignificant). Orange asterisks indicate comparison between H357-S/SCC9-S (si NTC) to H357-S/SCC9-S (si *POLK*) respectively (**i**) and (**ii**).

**Supplementary Figure 2:** **(A)** Bar graph showed the fold change in expression of γH2AX normalized to actin in H357-S, H357-R and H357-R *POLK* KO cells without and with cisplatin treatment, mean ± SEM (n=2). Statistical significance as determined by Ordinary one-way ANOVA, Dunnett's multiple comparisons test, (***p < 0.001, and ns- nonsignificant). **(B)** Bar graph depicted a fold change in expression of cleaved PARP1 normalized to actin in H357-S, H357-R and H357-R *POLK* KO cells without and with cisplatin treatment, mean ± SEM (n=2). Statistical significance as determined by Ordinary one-way ANOVA, Dunnett's multiple comparisons test, (*p < 0.05, ***p < 0.001, and ns- nonsignificant). **(C)** Representative cell cycle profiles of H357-S, H357-R and H357-R *POLK* KO cells at different time points after release from double thymidine block and cisplatin treatment were shown. G1, S, and G2M phases were demarcated (**i**). The percentage of cells in G1, S, and G2-M phases at 6 hours are estimated and plotted (**ii**). Data was represented as mean ± SEM (n=3). Statistical significance was determined by Ordinary one-way ANOVA, Tukey's multiple comparisons test, (*p < 0.05, and ns- nonsignificant). **(D)** Representative immunofluorescence staining images of H357-S, H357-R and H357-R *POLK* KO cells showed the expression of Ki-67 (red) in nucleus (DAPI, blue), and their corresponding merged images. Zoom-in merged view of dashed box area showed in upper right corner of corresponding image (**i**). Bar diagram showed the percentage of Ki-67 positive nuclei. Percentage of Ki-67 positive nuclei were calculated as per the formula described in materials and methods section (**ii**). Data was represented as mean ± SEM (n=3). Statistical significance was determined by Ordinary one-way ANOVA, Tukey's multiple comparisons test, (*p < 0.05, and ns- nonsignificant). Scale bars indicate 10 μm.

**Supplementary Figure 3: (A)** Co-immunoprecipitation was performed in H357-R whole cell extract using Polθ antibody. Interaction between Polθ with Polκ with was analyzed by Western blotting. IgG was considered as a negative control. The experiment was repeated twice. **(B)** Immunofluorescence co-localization analysis of H357-R cells showing Polθ (red), Polκ (green), DAPI (blue), and corresponding merged images. Merged images represented co-localization of Polθ and Polκ in the nucleus. Zoom-in merged (only green and red) view of dashed box area showed in upper right corner of corresponding image. Scale bars indicate 7.5 μm. **(C)** Western blot analysis shows ectopic expression of GFP-tagged WT, PIP mutant and Catalytically Dead (CD) mutants in H357-S and SCC9-S cells. β-actin was considered as an internal control (**i**) and (**ii**). **(D)** H357-S/ SCC9-S and their respective mutants were treated with the indicated concentrations of cisplatin for 48 hours and then an MTT assay was performed. These graphs indicate the mean ± SEM (n=3), two-way ANOVA, Tukey's multiple comparisons test (*p < 0.05, **p < 0.01, ***p < 0.001, ****p < 0.0001, and ns- nonsignificant) (**i**) and (**ii**). Purple asterisks indicate comparison between H357-R/SCC9-R to H357-S/SCC9-S cells respectively. Cyan, burgundy, and orange asterisks indicate comparison between H357-S/SCC9-S to their respective GFP-WT, GFP-PIP and GFP-CD overexpressed mutants.

**Supplementary Figure 4: (A)** Bar graph showed fold change in the expression of ATR, ATM, P-ATR, P-ATM, Chk1, Chk2, P-Chk1 and P-Chk2 normalized to actin in the cellular lysate of H357-S, H357-R and H357-R *POLK* KO cells without and with cisplatin treatment, mean ± SEM (n=2). Statistical significance as determined by two-way ANOVA, Dunnett's multiple comparisons test, (*p < 0.05, **p < 0.01, ***p < 0.001, and ****p < 0.0001). **(B)** Bar graph depicted fold change in the expression of MRE11, RAD50, NBS1, P-NBS1, CtIP, BRCA1, BRCA2, RAD54 and Rad51 normalized to actin in the cell free lysate of H357-S, H357-R and H357-R *POLK* KO cells without and with cisplatin treatment, mean ± SEM (n=2). Statistical significance as determined by two-way ANOVA, Dunnett's multiple comparisons test, (*p < 0.05, and ****p < 0.0001). **(C)** Bar graph showed fold change in the expression of KU70, KU80, DNA PKCs, P-DNA PKCs, Artemis, DNA LIG IV, XLF and XRCC4 normalized to actin in the cellular extract of H357-S, H357-R and H357-R *POLK* KO cells without and with cisplatin treatment, mean ± SEM (n=2). Statistical significance as determined by two-way ANOVA, Dunnett's multiple comparisons test, (*p <

0.05, **p < 0.01, ***p < 0.001, and ****p < 0.0001). **(D)** Bar graph showed fold change in the expression of Polκ, Chk1, Chk2, P-Chk1, P-Chk2, BRCA2, CtIP, Rad51, Artemis and P-NBS1 in nuclear extract of in H357-S, H357-R and H357-R *POLK* KO cells without and with cisplatin treatment, normalized to Histone H3, mean ± SEM (n=2). Statistical significance as determined by two-way ANOVA, Dunnett's multiple comparisons test, (*p < 0.05, **p < 0.01, ***p < 0.001, and ****p < 0.0001).

**Supplementary Figure 5:** Representative immunofluorescence images of H357-R and *POLK* KO cells with and without treatment with 5 µM of cisplatin showed nucleus (DAPI, blue), Chk1 (green) and Chk2, P-Chk1, P-Chk2, BRCA2, Artemis (red) and their corresponding merged images. Quantification of foci numbers is indicated in a bar diagram as mean ± SEM (n=3). Statistical significance was determined by two-way ANOVA, Šídák's multiple comparisons test, (*p < 0.05, ***p < 0.001, ****p < 0.0001, and ns-nonsignificant). Scale bars indicate 7.5 µm.

**Supplementary Figure 6: Polκ is also involved in cisplatin resistance of SCC9 cells. (A)** SCC9-R cell lines were transfected with NTC siRNA and *POLK* siRNA as described in Methods. For cisplatin treated sets cisplatin was given for 24 hours. Lysates were collected from indicated cells and Western blotting was performed with indicated antibodies. β-actin was used as an internal control. Representative best immunoblots from two independent experiments are shown. Densitometric fold change relative to proteins in SCC9-R cells has been provided just below each band. (**B**) Representative immunofluorescence images of SCC9-R and SCC9-R *POLK* knock-down cells for DAPI (blue) and Polκ (green) puncta, and corresponding merged images are shown. Quantification of Polκ foci numbers is indicated in a bar diagram as mean ± SEM (n=3). Statistical significance determined by Ordinary one-way ANOVA, Tukey’s multiple comparisons test, (****p < 0.0001, and ns- nonsignificant). Scale bars indicate 7.5 µm. **(C)** Representative alkaline comet images show DNA tailing in cells following treatment with 5 µM of cisplatin for 48 hours. The bar graph represents percentage of tail DNA. Data represented as (mean ± SEM; n=3). Statistical significance determined by two-way ANOVA with Tukey’s multiple comparisons test (**p < 0.01, ****p < 0.0001, and ns- nonsignificant). **(D)** Cells were treated with the indicated concentrations of cisplatin drug for 48 hours and then an MTT assay was performed. The graph indicates the mean ± SEM (n=3), Statistical significance determined by two-way ANOVA, Šídák's multiple comparisons test, (**p < 0.01, ****p < 0.0001, and ns- nonsignificant). Pink asterisks indicate comparison between SCC9-R (si NTC) to SCC9-R (si *POLK*). **(E)** Representative immunofluorescence images of SCC9-R and SCC9-R *POLK* knock-down cells show γH2AX (red) puncta and DAPI (blue), and corresponding merged images. Quantification of γH2AX foci numbers is indicated in a bar diagram as mean ± SEM (n=3). Statistical significance determined by Ordinary one-way ANOVA, Tukey’s multiple comparisons test, (**p < 0.01, ***p < 0.001, ****p < 0.0001, and ns-non significant). Scale bars indicate 7.5 µm.

**Supplementary Figure 7:** (**A**) Quantification of foci numbers in Fig. 8C is indicated in a bar diagram as mean ± SEM (n=3). Statistical significance was determined by two-way ANOVA, Šídák's multiple comparisons test, (****$p < 0.0001$, and ns- nonsignificant). (**B**) Co-immunoprecipitation was performed in H357-R whole cell extract using USP2 antibody. Interaction between USP2 with Polκ with was analyzed by Western blotting. IgG was considered as a negative control. The experiment was repeated twice.

**Supplementary Table 1:** Fork speed of IdU tract (μm/min) after replication restart in H357-S, H357-R and H357-R *POLK* KO cells as determined by DNA fiber assay (n=3) (**a**), Percentage of various DNA replication events in H357-S, H357-R and H357-R *POLK* KO cells after HU treatment as determined by DNA fiber assay (n=3) (**b**).

**Supplementary Table 2:** DNA replication fork speed (μm/min) without and with cisplatin treatment in H357-S, H357-R and H357-R *POLK* KO cells (n=3) (**a**), Percentage of various DNA replication events in H357-S, H357-R and H357-R *POLK* KO cells without and with treatment of cisplatin treatment as determined by DNA fiber assay (n=3) (**b**).

**Supplementary Table 3:** DNA replication fork speed (μm/min) in wild type and H357-R *POLK* KO cells harbouring various expression constructs (WT, PIP mutant and catalytically dead mutant) without and with cisplatin treatment as determined by DNA fiber assay (n=3) (**a**), Percentage of various DNA replication events of the same (n=3) (**b**).

**Supplementary Table 4:** List of dysregulated genes that are involved in de-ubiquitination pathway (**a**), and ubiquitin-proteasomal pathway have been given (**b**). Up-regulated and down-regulated genes were highlighted in different colours (green and orange) respectively.

**Supplementary Table 5:** DNA replication fork speed (μm/min) in wild type and H357-R *POLK* KO cells harbouring EV and UBZ-mutant construct without and with cisplatin treatment as determined by DNA fiber assay (n=3) (**a**), Percentage of various DNA replication events of the same (n=3) (**b**).

**Supplementary Table 6:** DNA replication fork speed (μm/min) without and with cisplatin treatment in H357-R (siNTC) and H357-R (si*USP18*) cells (n=3) (**a**), Percentage of various DNA replication events in H357-R (siNTC) and H357-R (si*USP18*) cells without and with treatment of cisplatin treatment as determined by DNA fiber assay (n=3) (**b**).
