## Supplementary figures and images for "PCNA-Polκ-Polδ/USP18 axes stabilize replication fork and restart to reduce cisplatin cytotoxicity"

### Supp. Figs 1-7

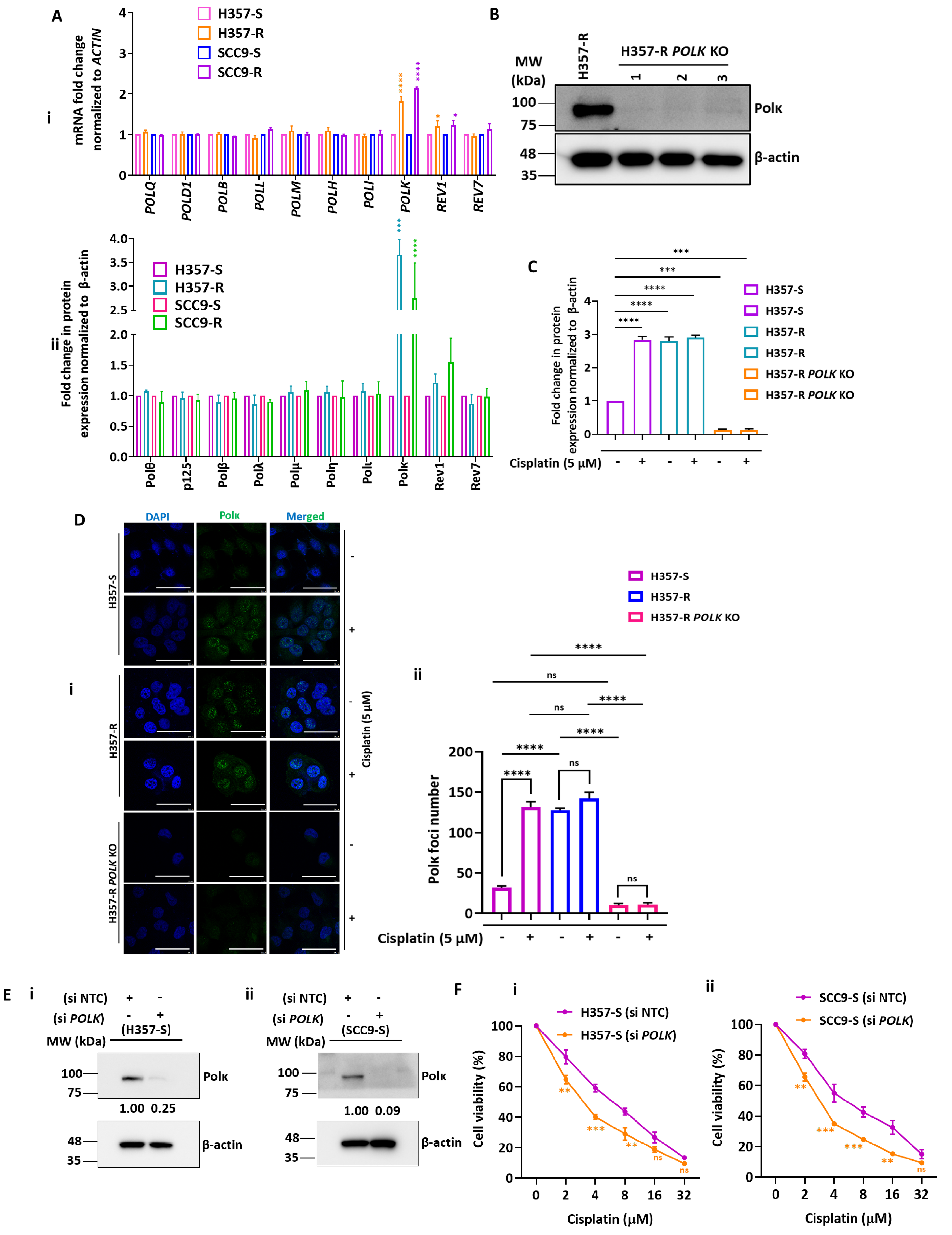


**Supp. Figure 1**

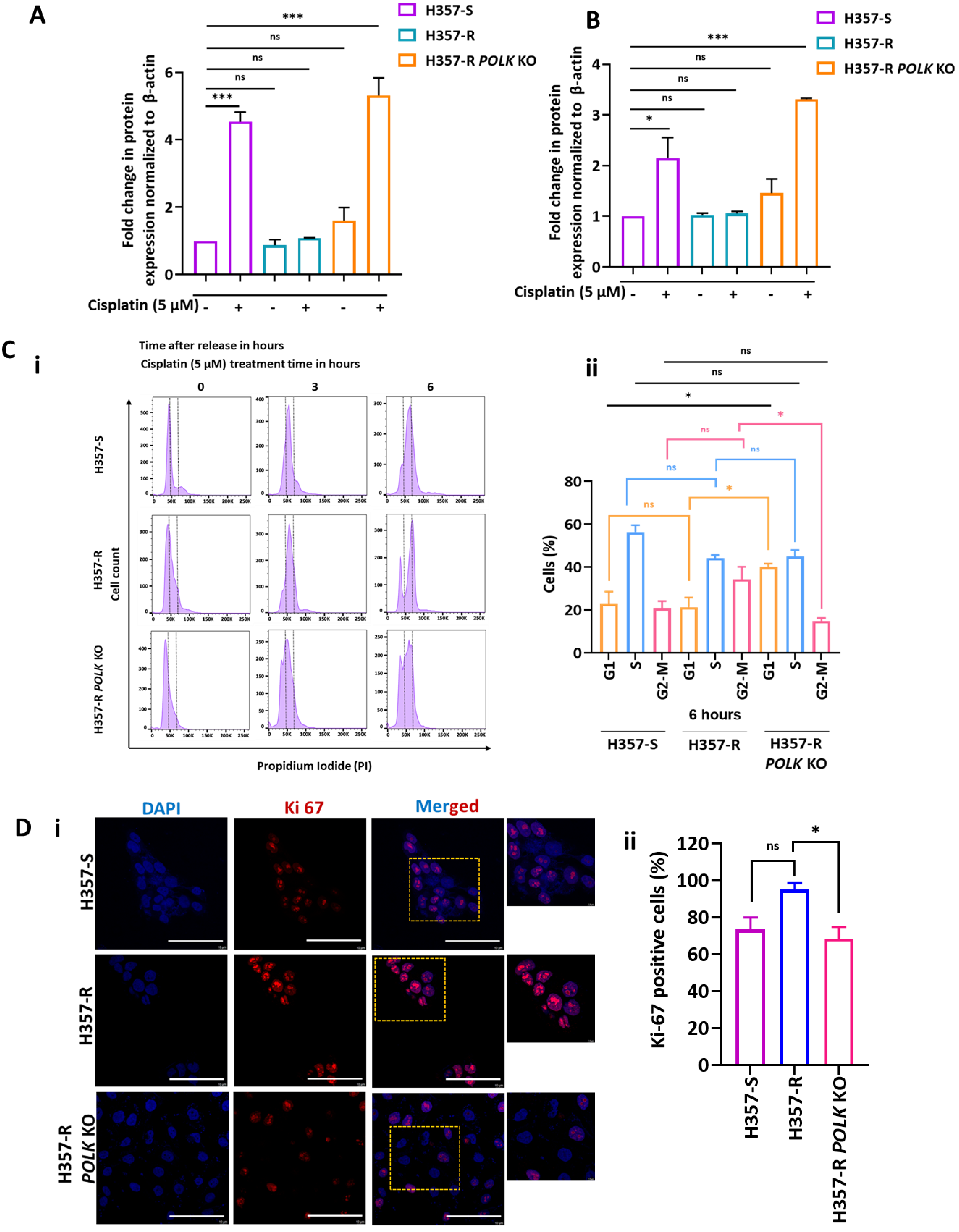


Supp. Figure 2

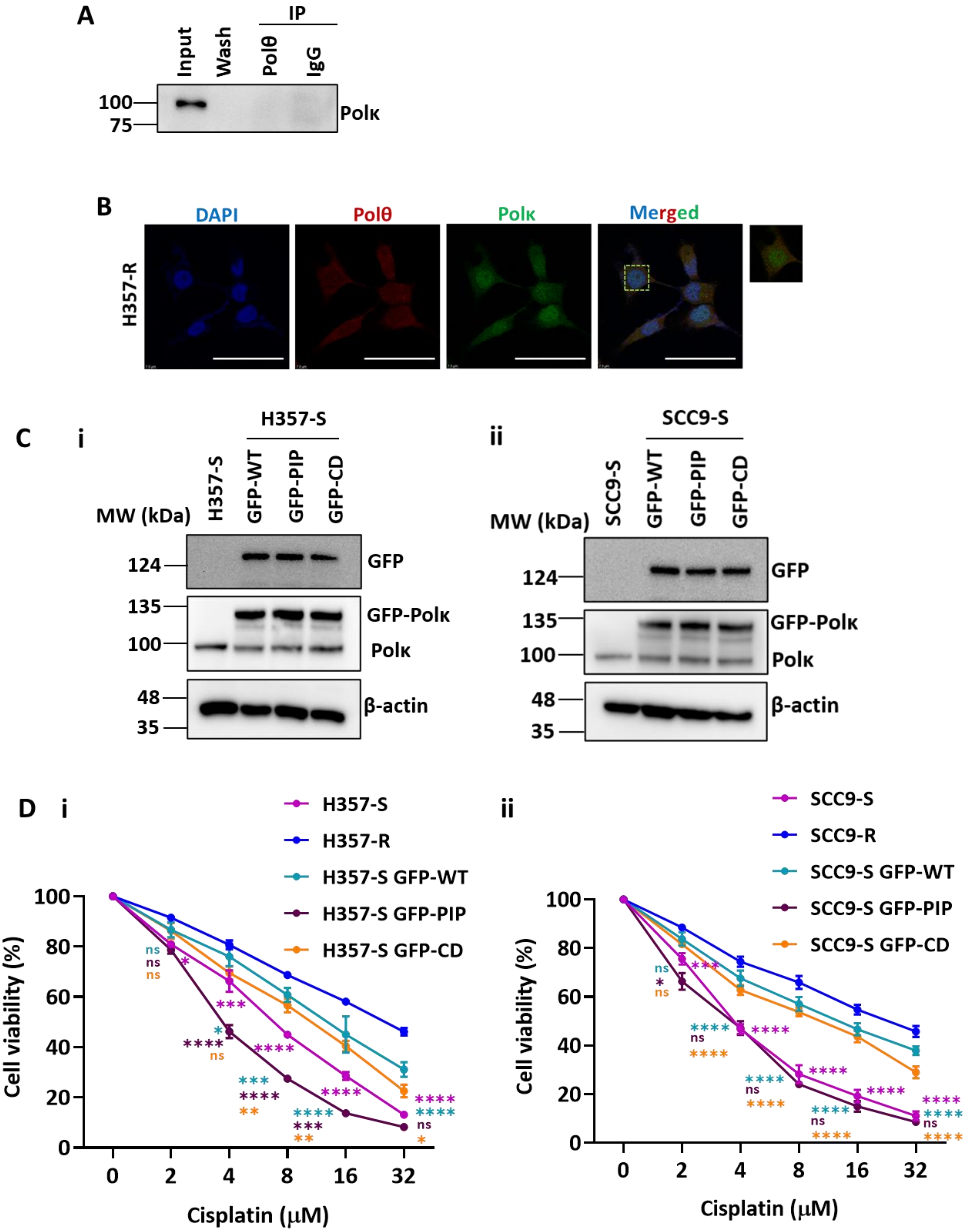


Supp. Figure 3

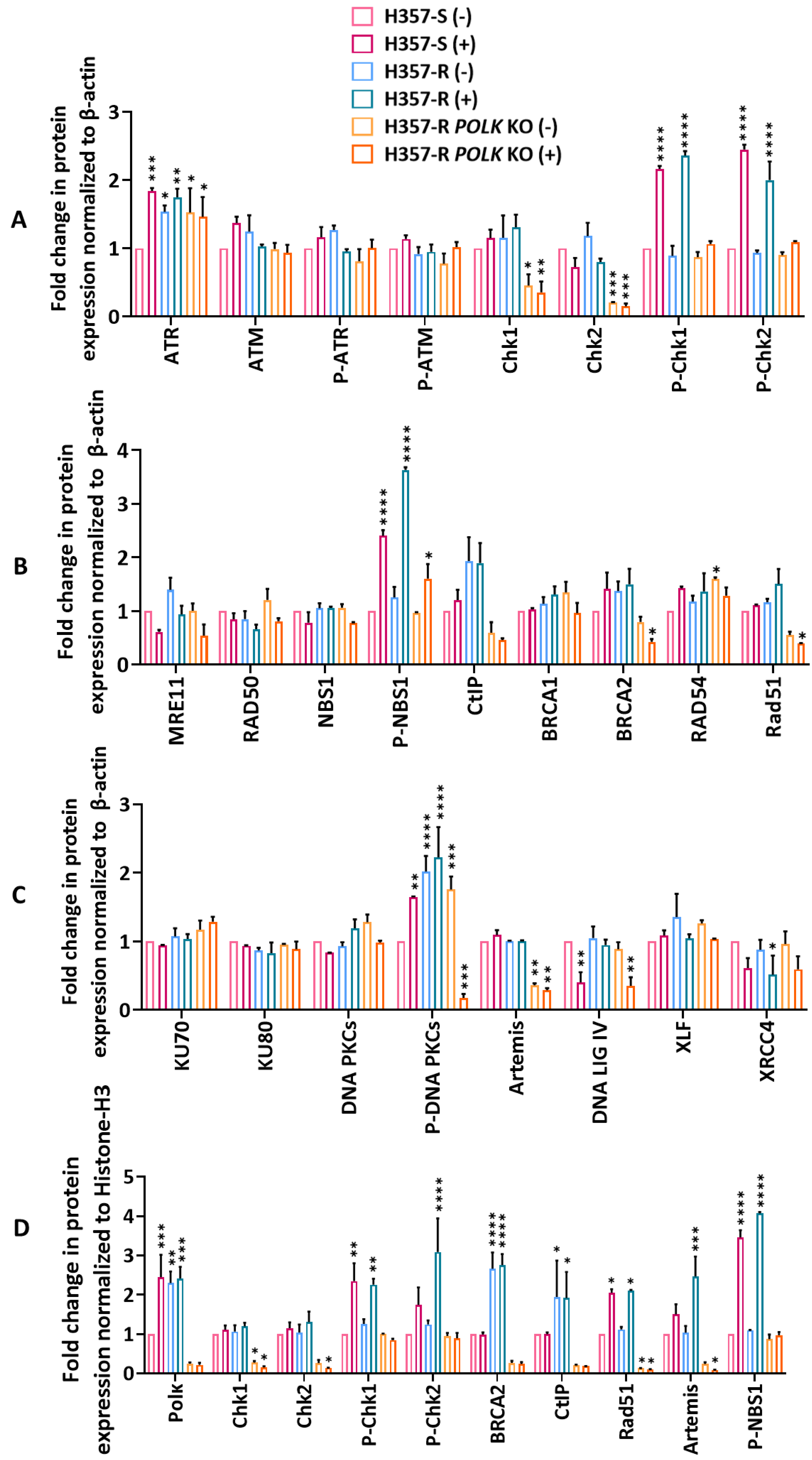


**Supp. Figure 4**

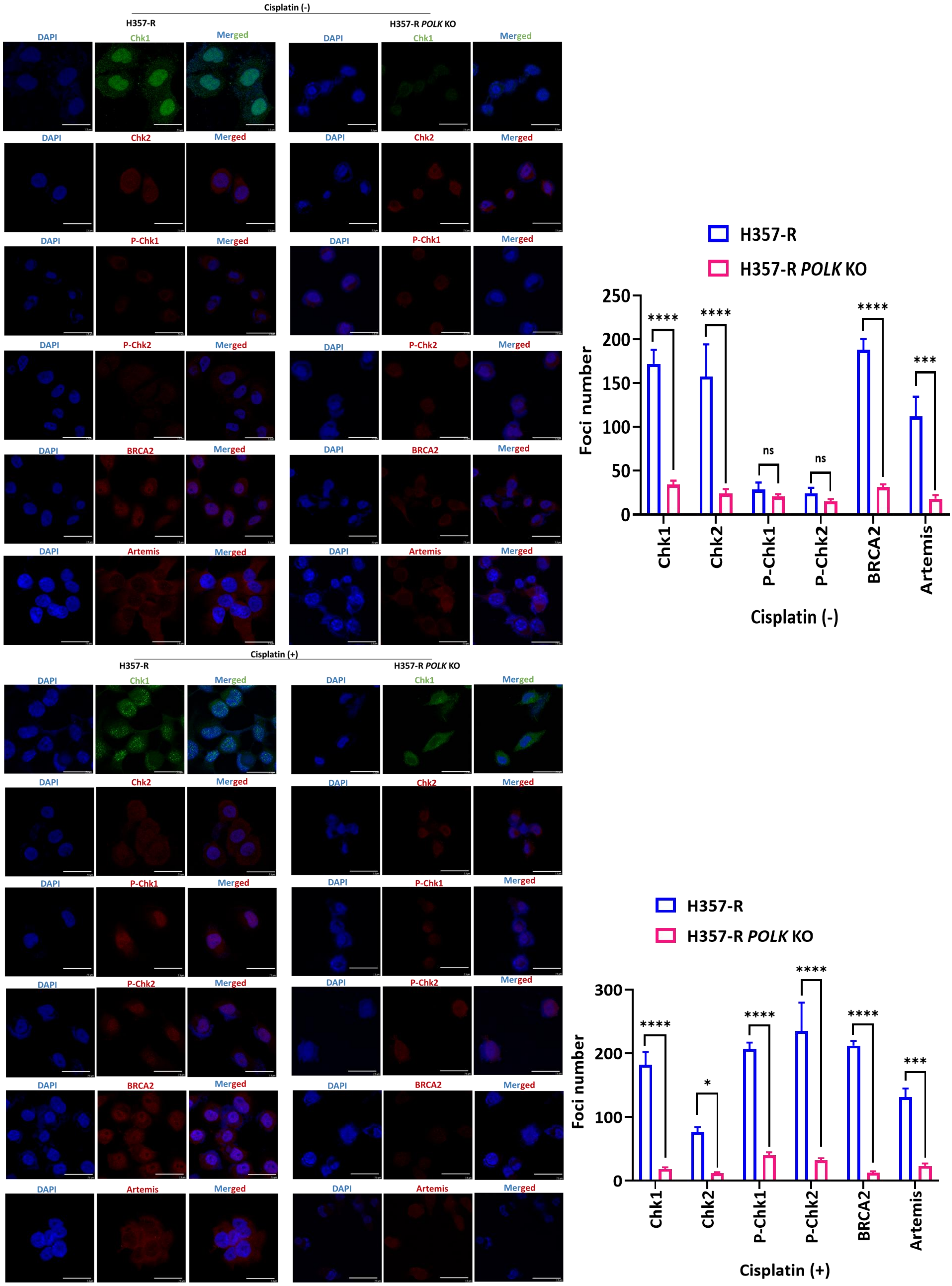


**Supp. Figure 5**

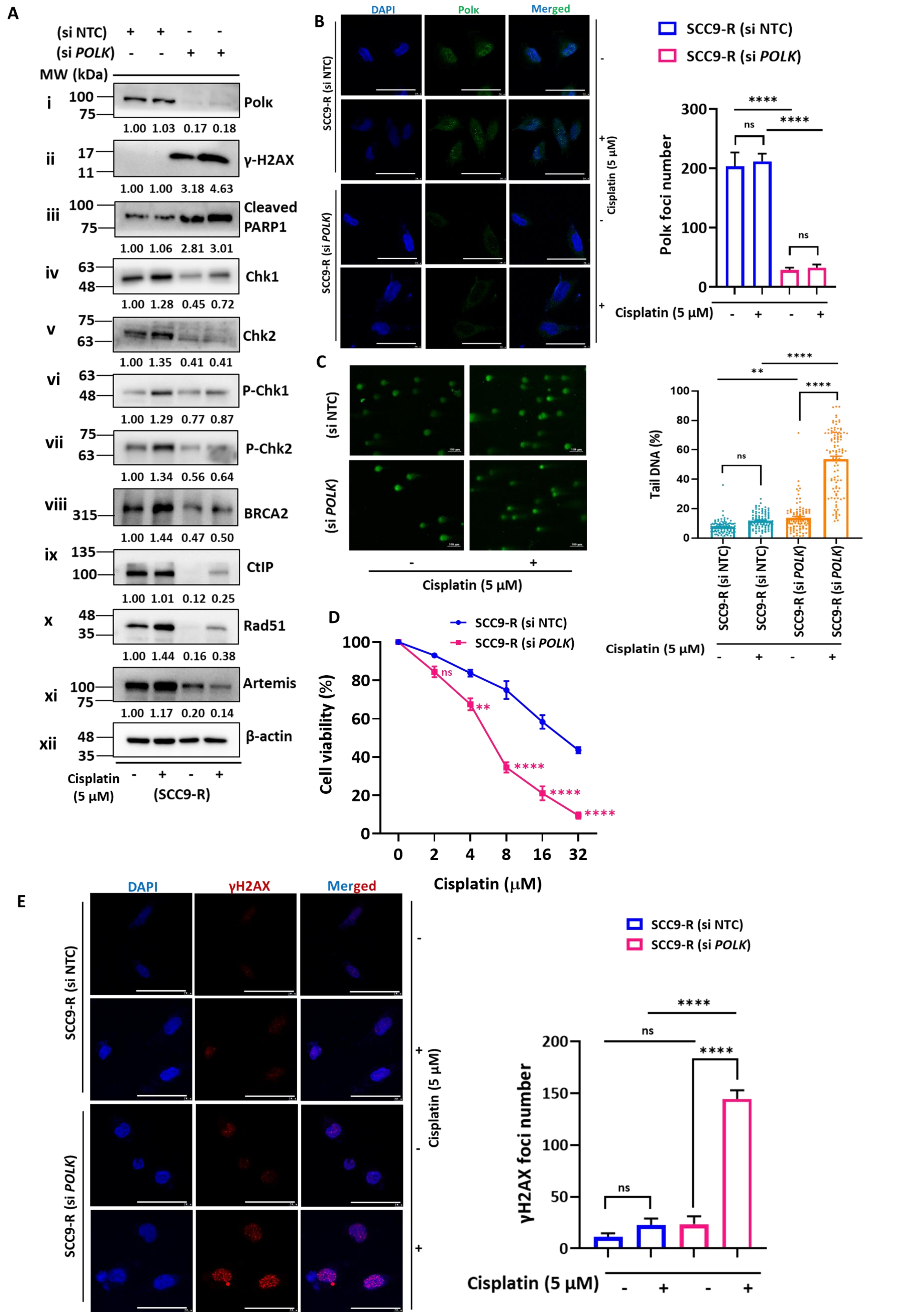


Supp. Figure 6

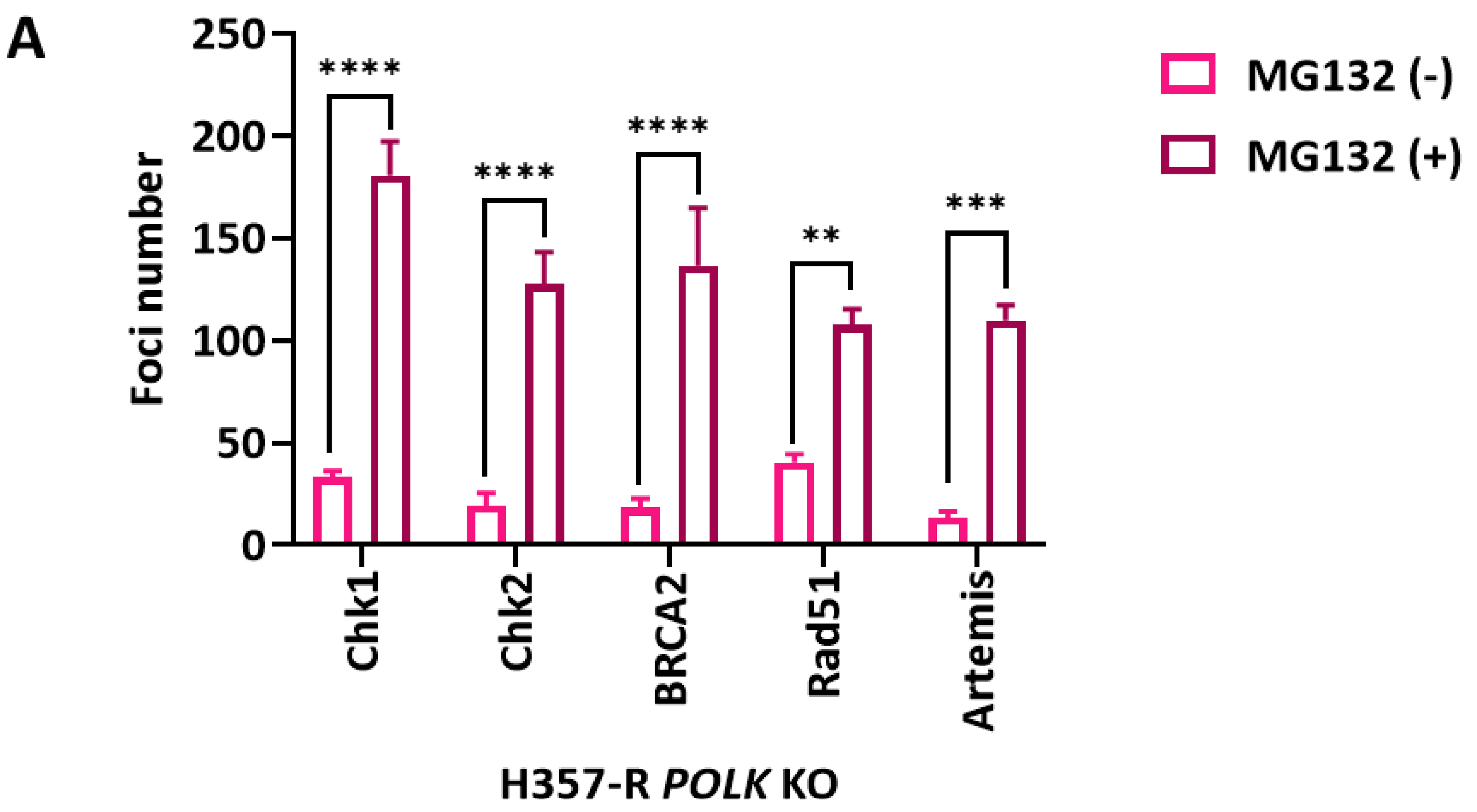


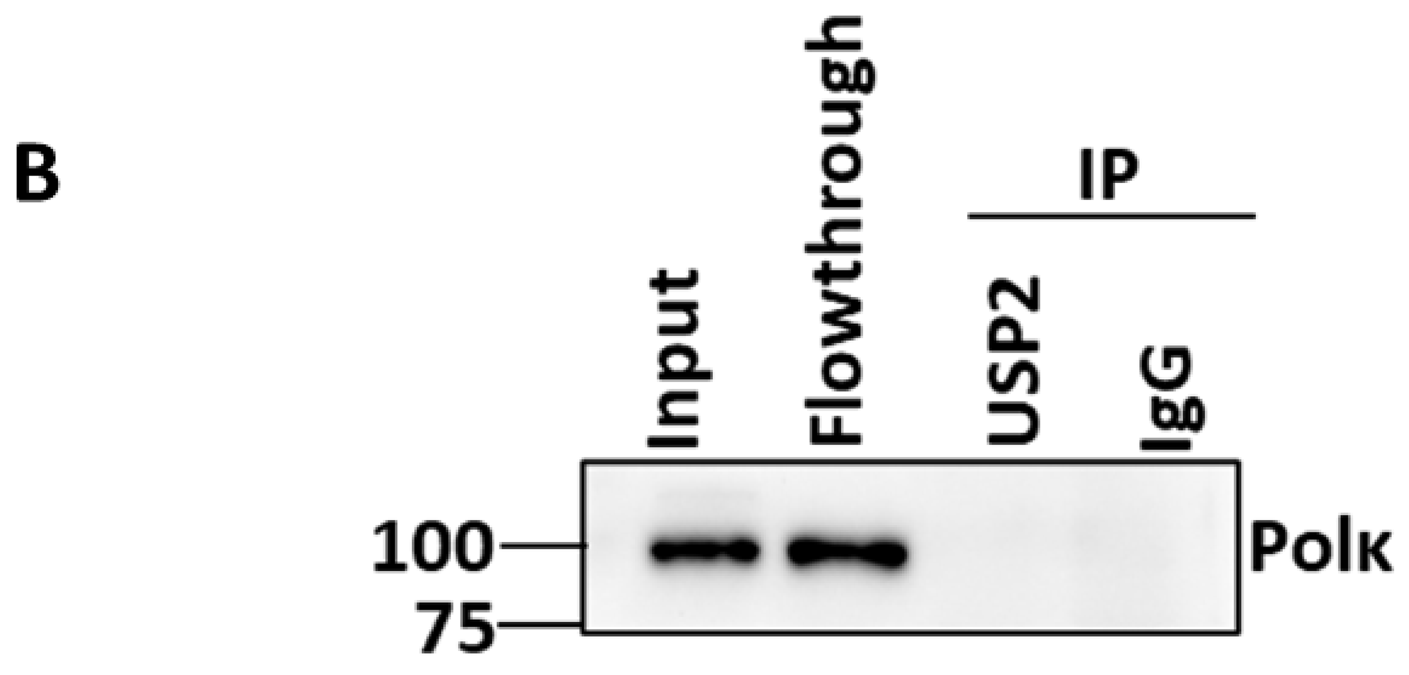


Supp. Figure 7
